# The human SKI complex prevents DNA-RNA hybrid-associated telomere instability

**DOI:** 10.1101/2020.05.20.107144

**Authors:** Emilia Herrera-Moyano, Rosa Maria Porreca, Lepakshi Ranjha, Eleni Skourti, Roser Gonzalez-Franco, Ying Sun, Emmanouil Stylianakis, Alex Montoya, Holger Kramer, Jean-Baptiste Vannier

## Abstract

Super killer (SKI) complex is a well-known cytoplasmic 3′ to 5′ mRNA decay complex that functions with the exosome to degrade excessive and aberrant mRNAs. Recently, SKIV2L, the 3′ to 5′ RNA helicase of the human SKI (hSKI) complex has been implicated in the degradation of nuclear non-coding RNAs escaping to the cytoplasm. Here, we show that hSKI is present in the nucleus, on chromatin and in particular at telomeres during the G2 cell cycle phase. In cells, SKIV2L prevents telomeric loss, and DNA damage response activation, and its absence leads to DNA-RNA hybrid-mediated telomere fragility. Moreover, we demonstrate that purified hSKI complex preferentially unwinds telomeric DNA-RNA hybrids *in vitro*. Taken together, our results provide a nuclear function of the hSKI complex in overcoming replication stress caused by aberrant processing of telomeric DNA-RNA hybrids and thus maintaining telomere stability.

## Introduction

To tackle the challenge of aberrant or excessive cytoplasmic and nuclear RNA molecules, cells have evolved different RNA decay pathways. Among these, the nonsense-mediated mRNA decay (NMD) surveillance pathway is involved in the degradation of mRNAs presenting premature translation termination (Isken and Maquat 2007) and also works as a quality-control system regulating the expression of physiological RNAs (Hug et al. 2016). Although human NMD has been widely investigated for its cytoplasmic functions (Singh et al. 2007), some of its core components have been identified in the nucleus, targeting several essential biological processes including telomere homeostasis (Azzalin and Lingner 2006; Azzalin et al. 2007).

Telomeres are transcribed by RNA polymerase II from subtelomeric regions towards the chromosome ends into the long non-coding telomeric repeat-containing RNA (TERRA) (Azzalin et al. 2007; Schoeftner and Blasco 2008; Azzalin and Lingner 2015). TERRA transcription and its association with telomeres is co-ordinated throughout the cell cycle to allow for correct post-replication processing of telomeres but also to avoid collisions between DNA replication and transcription (Porro et al. 2010; Maicher et al. 2012). Due to the complementary nature of TERRA to telomeric DNA, R-loops, constituted by the DNA-RNA hybrid and the displaced DNA strand, may form at telomeres and result in telomeric DNA replication stress while they also act as key mediators of telomere length dynamics (Balk et al. 2013; Arora et al. 2014; Graf et al. 2017; Sagie et al. 2017). TERRA levels vary in a cell cycle-dependent manner, peaking in G1 phase, decreasing in S phase and reaching the lowest level in S/G2 in HeLa cells (Porro et al. 2010), which suggests the presence of a regulatory mechanism. Indeed, the RNA helicase UPF1, essential for the NMD pathway, binds to chromatin during S phase (Azzalin and Lingner 2006) and particularly to telomeres *in vivo* (Azzalin et al. 2007). Its activity is essential to ensure replication and telomere length homeostasis by mediating the displacement of TERRA from telomeres (Chawla et al. 2011).

In *Saccharomyces cerevisiae*, the cytoplasmic 3′-5′ mRNA decay relies on the super killer (SKI) and exosome complexes. The yeast SKI tetramer is composed of Ski2, Ski3 and two subunits of Ski8 (SKIV2L, TTC37, WDR61 in human, respectively) (Halbach et al. 2013). Ski2 presents a 3’ to 5’ RNA helicase activity that channels the RNA into the exosome for further degradation via interaction with Ski7 (Wang et al. 2005; Halbach et al. 2013). Consistent with the function of its yeast homolog, human SKIV2L facilitates cytoplasmic degradation of mRNAs and viral RNA by the exosome and plays a role in RNA interference (Chen et al. 2001; van Hoof et al. 2002; Orban and Izaurralde 2005; Aly et al. 2016). Recently, the human SKI (hSKI) complex was shown to extract mRNA by directly interacting with the ribosomal complexes *in vitro* (Zinoviev et al. 2020). Indeed, SKIV2L works in a translation surveillance programme with the RNA binding protein AVEN, preventing ribosome stalling by eliminating RNA transcripts (Tuck et al. 2020). The implication of hSKI in different RNA regulatory pathways and its localisation in both the cytoplasm and the nucleus (Qu et al. 1998; Zhu et al. 2005), suggests a possible role for the hSKI complex in nuclear RNA surveillance.

Our data show that the hSKI complex also localises to chromatin and is enriched at telomeres during the G2 phase of the cell cycle. Purified hSKI not only interacts with RNA but also has a high affinity for single-stranded telomeric DNA. Strikingly, hSKI binds DNA-RNA hybrids *in vitro* and its recruitment to telomeres is dependent on telomeric DNA-RNA hybrids. Loss of SKIV2L decreases the amount of telomeric DNA-RNA hybrids in G2 and provokes telomere abnormalities. These include ATM-mediated DNA damage response induction caused by increased telomere loss and fragility, with the latter being suppressed by the depletion of TERRA and by the over-expression of RNase H1. Overall, our results describe an unforeseen function of hSKI in the maintenance of telomere integrity before mitosis through the regulation of telomeric DNA-RNA hybrids homeostasis.

## Results

### The human SKI (hSKI) complex is recruited to telomeres, particularly in G2

In order to assess the localisation of hSKI within HeLa cells, we firstly employed cell fractionation. The three components of hSKI: SKIV2L, TTC37 and WDR61 were detected in the whole cell lysate and the chromatin-bound fraction, while the soluble fraction only contained SKIV2L (Figure 1A). Because other factors of the nonsense-mediated mRNA decay (NMD) pathway have been associated with telomeres in S phase (Chawla et al. 2011), we assessed the binding of hSKI components to telomeres throughout the cell cycle. Upon cell synchronisation (Figures S1A and S1B), telomere binding was examined using Proteomic of Isolated Chromatin segments (PICh) (Dejardin and Kingston 2009; Porreca et al. 2020). The PICh analysis was conducted using unique peptides identified in each condition and the respective label-free quantification values. As expected, peptides from the Shelterin factors: TRF2, TRF1, RAP1, TIN2, TPP1 and POT1 were found enriched in the reads obtained with the telomeric probe compared to the scrambled probe, which confirms specificity of the experiment (Figure S1C). Interestingly, the three components of the hSKI complex were also found in the telomere probe-derived peptide reads, with SKIV2L and TTC37 significantly enriched during G2 phase (Figure 1B).

**Figure 1.**
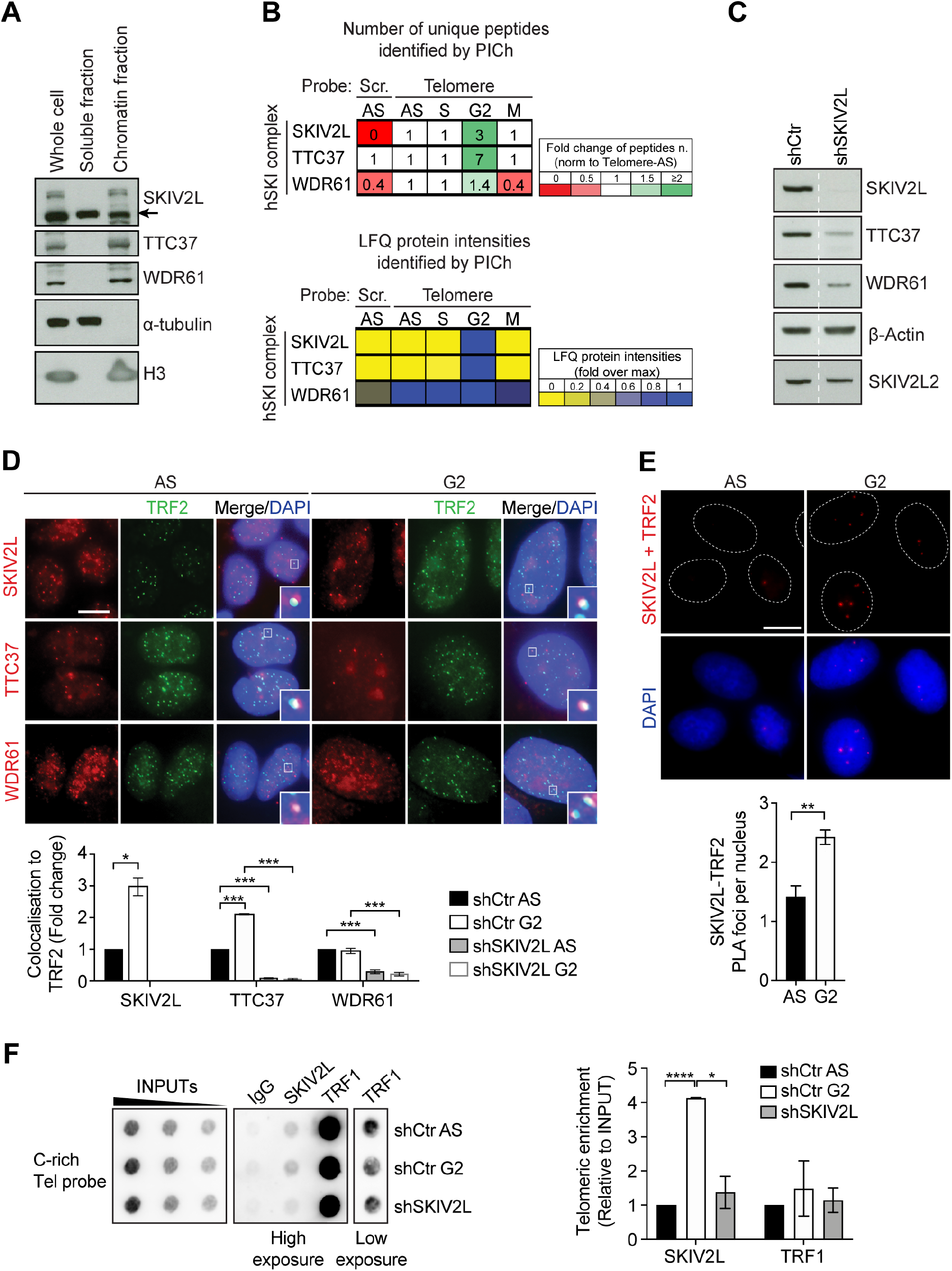
Human SKI complex binds telomeres and is enriched in G2. (A) Subcellular fractionation assay in asynchronous HeLa cells. (B) Proteomics of isolated chromatin segments analysis showing the binding of hSKI to telomeres throughout the cell cycle. (C) WB of hSKI (SKIV2L, TTC37 and WDR61) and SKIV2L2 in shCtr (Control) or shSKIV2L cells. (D) Immunofluorescence showing co-localisation of hSKI with TRF2 in asynchronous (AS) and G2-synchronised cells. % of total number of SKIV2L, TTC37 or WDR61 foci colocalising with TRF2 are normalised to shCtr AS samples (means ± SEM, n = 2 independent experiments, scale bar 15 µm). t test *p<0.05, ***p <0.001. (E) Proximity ligation assay of SKIV2L-TRF2, showing increasing number of foci in G2 synchronised cells (means ± SEM, n = 3-4 independent experiments, at least 600 cells scored per condition, scale bar 10 µm). t test **p <0.01. (F) ChIP-dot blot of SKIV2L and TRF1 in AS and G2-synchronised shCtr and AS shSKIV2L cells (means ± SEM, n >= 2). t test *p<0.05, ****p <0.0001.

To further validate the presence of the hSKI complex at telomeres, we successfully generated stable SKIV2L knockdown (shSKIV2L) and control (shCtr) HeLa cells using short hairpin RNAs (Figure 1C). Notably, the depletion of SKIV2L was accompanied by decreased protein levels of the other two hSKI components (TTC37 and WDR61) but had no effect on the protein expression of its paralog, SKIV2L2, known to regulate telomerase RNA levels (Nguyen et al. 2015). Therefore, any phenotype described herein upon SKIV2L depletion is SKIV2L2-independent. Using immunofluorescence, we examined the co-localisation of each of the hSKI components with TRF2, a telomeric protein. All three hSKI components were detected in the nucleus, and at telomeres in both asynchronous and G2 cells (Figure 1D). Notably, the localisation of both SKIV2L and TTC37 at telomeres was significantly increased in G2 cells, consistent with the PICh data. In agreement with the reduced protein levels upon SKIV2L depletion (Figure 1C), we observed loss of co-localisation of all three hSKI components with TRF2, both in asynchronous and G2 cells (Figures 1D and S1D), which suggests that hSKI is recruited as a complex at telomeres. Similarly, we observed localisation of SKIV2L to telomeres using SKIV2L-immunofluorescence coupled with telomeric FISH. The increase of SKIV2L at telomeres in G2-synchronised cells was also abolished upon SKIV2L depletion (Figure S1E). In addition, we confirmed the enhanced SKIV2L-TRF2 co-localisation in G2 phase by proximity ligation assay (PLA) (Figures 1E and S1F). PLA with individual antibody (TRF2 or SKIV2L) and together in the knock-down cell line was used to ensure specificity of the results (Figure S1F). Finally, telomeric ChIP-dot blot confirmed higher level of SKIV2L at telomeres in G2 cells, which was significantly reduced in SKIV2L-depleted cells (Figure 1F). Notably, we did not notice any alteration in the recruitment of TRF1 and TRF2 to telomeres, respectively by ChIP and IF (Figure 1F and S2A), and no changes in TRF1, TRF2 and TIN2 protein expression by western blot (Figure S2B), in cells deficient for SKIV2L. This suggests that SKIV2L downregulation has no major effect on the expression of some shelterin proteins.

Taken together, these results indicate that hSKI localises to telomeres, particularly during the G2 phase of the cell cycle.

### SKIV2L suppresses telomere loss, telomere fragility and DNA damage signalling activation

To shed light on the potential telomeric function of hSKI, we investigated whether the depletion of the hSKI components would perturb telomere stability. We used telomere quantitative-FISH analysis of metaphases of HeLa and HT1080-ST cells depleted for SKIV2L and TTC37 by siRNAs to look for telomere abnormalities (Figure 2A-2B and S2B-C). We noticed an increase in telomere fragility and telomere loss that are respective markers of replication stress and instability at telomeres (Sfeir et al. 2009; Vannier et al. 2012), in both cell lines depleted for SKIV2L and TTC37. Notably, the depletion of the third component of the complex, WDR61, did not induce significant telomere fragility or loss in HeLa and HT1080-ST cells (Figure S2C-E). The same phenotypes of telomere fragility and loss were reproduced in the stable shRNA HeLa cell line (short hairpin targeting a different region of SKIV2L mRNA compared to the siRNA pool) showing more than 2.5- and 3-fold increase, respectively, compared to the stable shCtr HeLa cell line (Figure 2C). To test whether these phenotypes are dependent on replication stress at telomeres, we treated the stable cell lines with low doses of the DNA polymerase inhibitor aphidicolin (APH). Telomere fragility caused by SKIV2L depletion is slightly more pronounced than in control cells treated with APH (used as a positive control), which confirms that SKIV2L is important for the maintenance of telomere integrity (Figure 2C). Surprisingly, SKIV2L-depleted cells treated with APH showed suppression of telomere fragility to background levels while telomere loss was unaltered, suggesting that the phenotype of telomere fragility is replication-dependent (Figure 2C). Indeed, APH-treated cells exhibit a delay in the progression through the S phase, while SKIV2L knockdown has no effect on the cell cycle profile of HeLa cells (Figure 2D).

**Figure 2.**
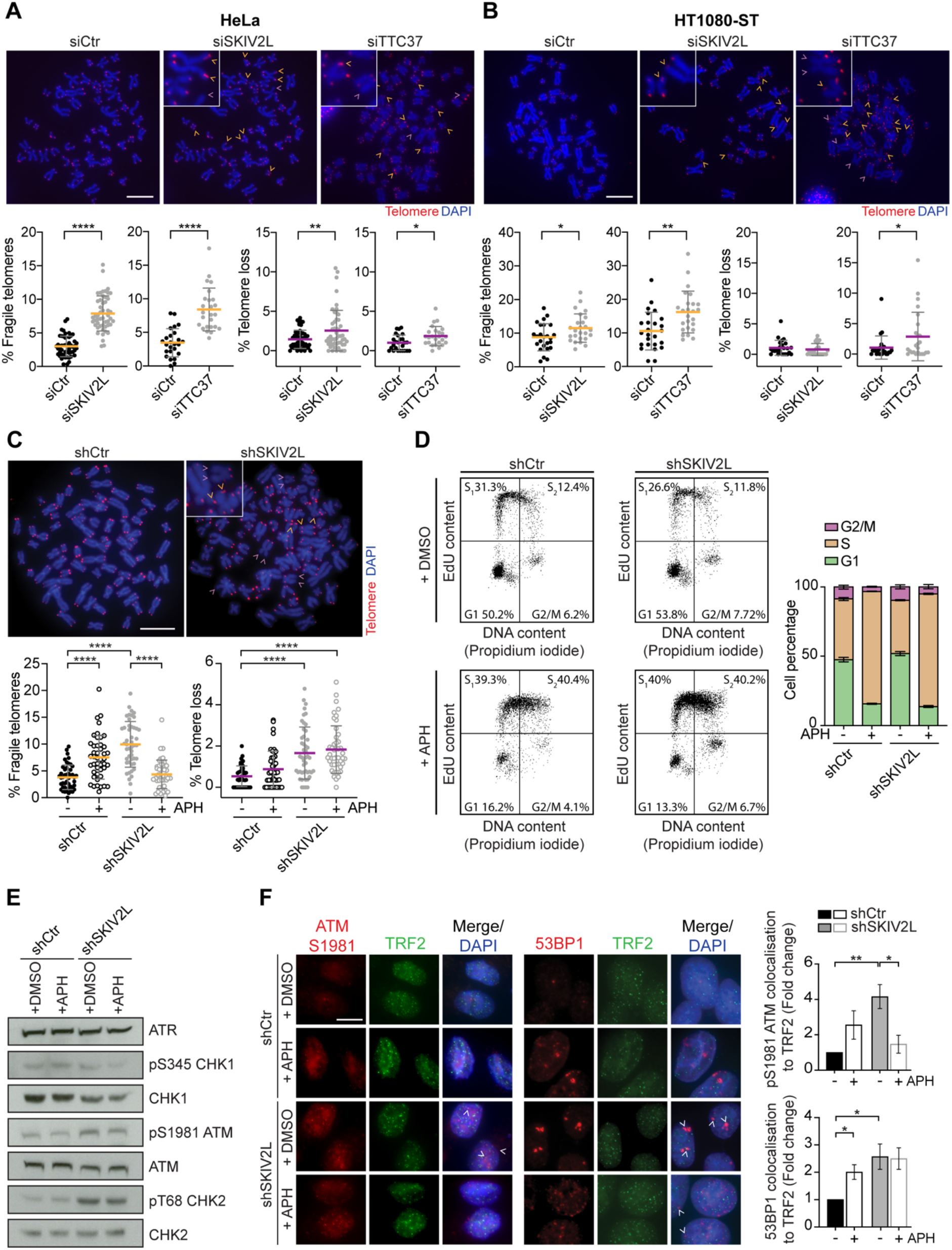
SKIV2L prevents telomere fragility, loss and DNA damage signalling activation. (A-B) Telomere FISH analysis in HeLa and HT1080-ST cells using siRNAs: % of telomere fragility (yellow) and loss (purple) per metaphase in siCtr, siSKIV2L and siTTC37 (means ± SD, n >= 25 metaphases, 1-2 independent experiments, scale bar 10 µm). t test *p<0.05, **p <0.01, **** p <0.0001. (C) Telomere FISH analysis in HeLa cells using shRNAs: shCtr and shSKIV2L treated with DMSO (−, control) or APH (+) (n >= 45 metaphases, 2 independent experiments). Details are as in (A). (D) Flow cytometry analysis of the cell cycle distribution in shCtr and shSKIV2L HeLa cells untreated (DMSO) or APH treated. DNA was stained with propidium iodide and cells were incubated with EdU to mark newly synthesised DNA. The percentage of cells in G1, early S phase (S_1_), late S phase (S_2_) and G2/M phases is indicated (n=10000 cells) (mean ± SEM, n = 3 independent experiments, data from S1 and S2 are merged as dataset S). (E) WB analysis of ATR, pS345 CHK1, CHK1 (loading control), pS1981 ATM, ATM (loading control), pT68 CHK2 and CHK2 (loading control) in shCtr and shSKIV2L HeLa cells treated with DMSO or APH. (F) TRF2, pS1981 ATM and 53BP1 IF staining in shCtr and shSKIV2L HeLa cells treated with DMSO (−) or APH (+). % of pS1981 ATM or 53BP1 foci colocalising with TRF2 are normalised to shCtr untreated samples (mean ± SEM, n = 3 independent experiments, at least 100 cells scored per experiment, scale bar 15 µm). t test * p<0.05, ** p<0.01.

Since we identified various telomere abnormalities in SKIV2L knockdown cells, we asked if this is accompanied with the activation of the DNA damage response. Even though no change in the pS345 phosphorylation levels of the ATR-effector kinase CHK1 was detected; we observed an increase in the levels of activated ATM (pS1981 ATM autophosphorylation), as well as activation of its effector kinase CHK2 (pT68 phosphorylation) in cells depleted of SKIV2L. No differences were observed between APH treated and untreated SKIV2L-deficient cells (Figure 2E).

Interestingly, when the DNA damage response was monitored specifically at telomeres via co-localisation with TRF2 (Figure 2F), a 4-fold increase in ATM activation was detected upon SKIV2L depletion. Hindering DNA replication with APH treatment in SKIV2L-deficient cells suppressed this activation, in agreement with the APH-dependent telomere fragility suppression. Furthermore, to measure the levels of DNA damage at telomeres, we quantified telomeric dysfunction-induced foci (TIFs) by the localisation of 53BP1 foci at telomeres (Takai et al. 2003). In both APH treated and SKIV2L-depleted cells, we observed increased levels of TIFs when compared to control cells (Figure 2F). However, this increase was not rescued by treatment of SKIV2L-depleted cells with APH (Figure 2F). Therefore, we conclude that telomere fragility and ATM activation are caused by DNA replication stress in SKIV2L-depleted cells, while telomere loss and 53BP1-dependent DNA damage occur independently of DNA replication (Figure 2C and 2F). In summary the above results confirm that SKIV2L is a regulator of telomere maintenance and its absence leads to abnormal processing of telomeres and DNA damage.

### Purified hSKI complex binds RNA and DNA substrates containing telomeric repeats

To elucidate the potential nuclear function of hSKI and understand how it interacts with telomeres, we resorted to its purification followed by *in vitro* biochemical assays. We designed N-terminus MBP-tagged SKIV2L, N-terminus His-Flag tagged TTC37, untagged WDR61 and purified from insect cells the hSKI complex by subsequent affinity and size exclusion chromatography (Figure 3A and 3B).

**Figure 3.**
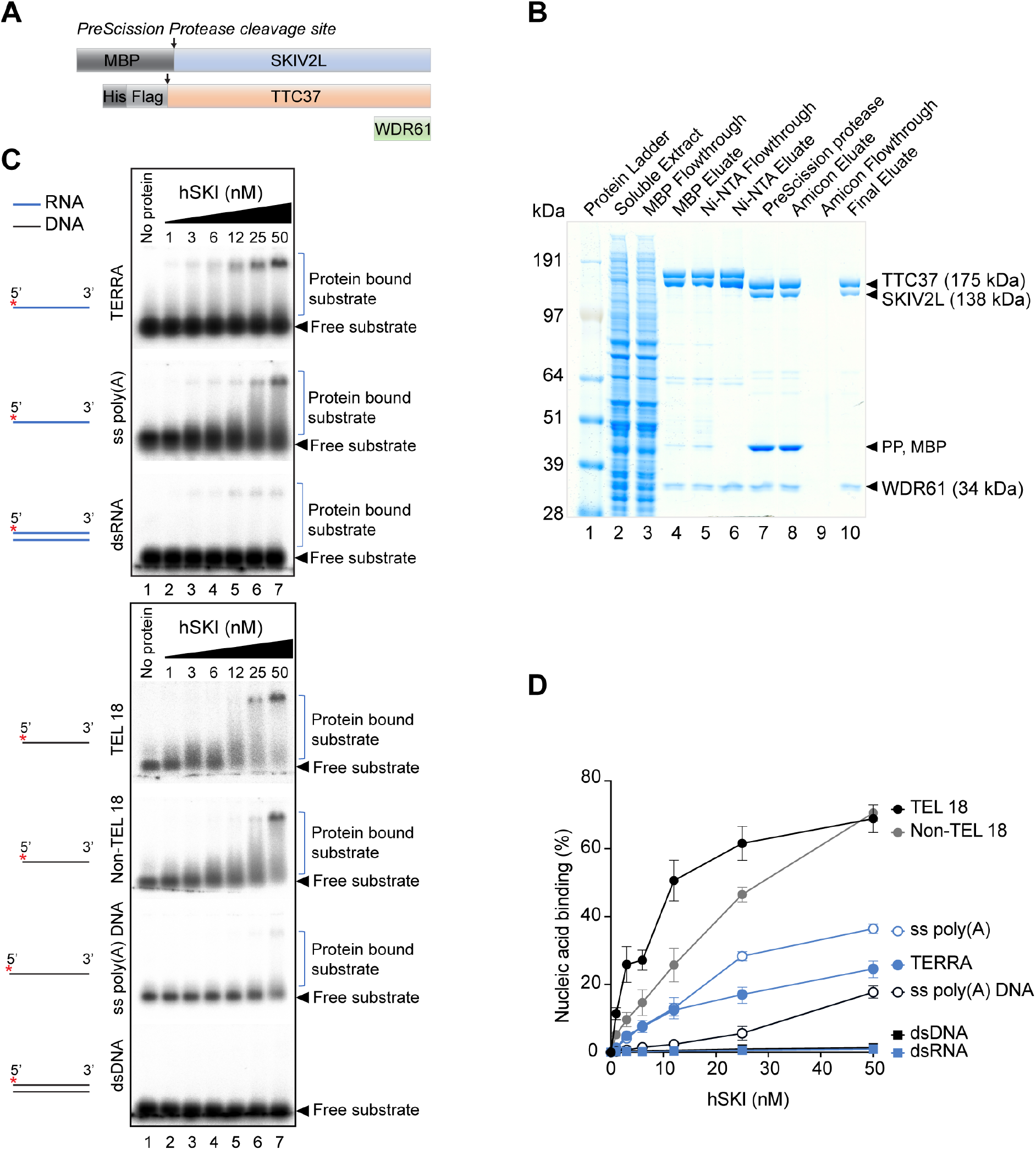
Purified recombinant hSKI binds preferentially telomeric ssDNA. (A) hSKI constructs. MBP, Maltose binding protein; His, 6x histidine; Flag, 3x flag. (B) Coomassie blue SDS-PAGE gel showing purified hSKI (lane 10) and different purification fractions. PP, PreScission protease. (C) Electrophoretic mobility shift assays showing binding of hSKI to different RNA and DNA substrates. Blue lines denote RNA, black lines denote DNA and asterisk indicates radioactive ^32^P label. (D) Quantification of (C) showing % of nucleic acid binding calculated as protein bound substrate signal relative to free substrate signal (mean ± SEM, n = 2-4).

As SKIV2L contains two putative RecA-like domains and interacts with mRNA, we set out to evaluate the binding properties of purified hSKI complex to various nucleic acids using electrophoretic mobility shift assays (Figure 3C). First, we found that similar to its yeast homolog, hSKI can bind single stranded RNA (ssRNA) containing a poly(A) tail but also ssDNA of similar sequence (ss poly(A) and ss poly(A) DNA, respectively) (Figures 3C and 3D). Since we observed that hSKI localises to telomeres, we then evaluated the binding affinity of hSKI to various telomeric and non-telomeric RNA and DNA substrates including ssRNA and ssDNA (TERRA, TEL 18 and non-TEL 18) and double stranded RNA and DNA (dsRNA/ dsDNA) (Figure 3C). Strikingly, we found that hSKI favours binding to telomeric ssDNA (TEL18, K_d_ = 12 nM) as compared to non-telomeric ssDNA (Non-TEL18, K_d_ = 30 nM) of similar length (Figures 3C and 3D). Binding to ssRNA of telomeric sequence, that mimics TERRA, was also observed; however, the affinity was lower than for telomeric ssDNA (TERRA *vs* TEL18) (Figures 3C and 3D). Importantly, no binding to blunt-ended dsDNA and dsRNA was observed. These results show that hSKI can bind single-stranded telomeric oligonucleotides.

### Purified hSKI binds telomeric DNA-RNA hybrids with single-stranded 3’ end

Previously, SKIV2L and TTC37 were identified among 400 factors enriched in a synthetic TERRA-interacting proteins screen using SILAC-based quantitative mass spectrometry (Scheibe et al. 2013). As hSKI was able to bind both ssDNA and ssRNA, we next tested the complex for its binding specificity towards various duplex substrates containing either 3’ or 5’ overhangs that could mimic telomere structures *in vitro* (Figure 4A and S3A). Consistent with its higher affinity for telomeric ssDNA rather than ssRNA, hSKI showed a preference for substrates containing telomeric DNA 3’ overhangs (3’ TEL D:D, K_d_ 6 nM) compared to substrates containing RNA 3’ overhangs (3’ TERRA R:R, K_d_ of 12 nM; 3’ TERRA R:D, K_d_ of 25 nM; Figure 4A). Strikingly, hSKI showed a greater preference for G-rich 3’ overhangs compared to non-telomeric 3’ overhang (Figure 4A, 3’ Non-TERRA-R:D) and poly(A) 3’ overhangs (Figure S3A). Furthermore, hSKI showed poor binding for structures containing 5’ overhang (5’ TEL D:D and 5’ Non-TERRA R:R) (Figure S3A). The data indicate that hSKI has preferential affinity for G-rich telomeric substrates harbouring 3’ DNA or RNA overhangs, thus advocating a function on telomere structures (G-overhang or DNA-RNA hybrids). Indeed, in a recent study, SKIV2L and TTC37 were found as factors pulled-down with synthetic DNA-RNA hybrids (Wang et al. 2018).

**Figure 4.**
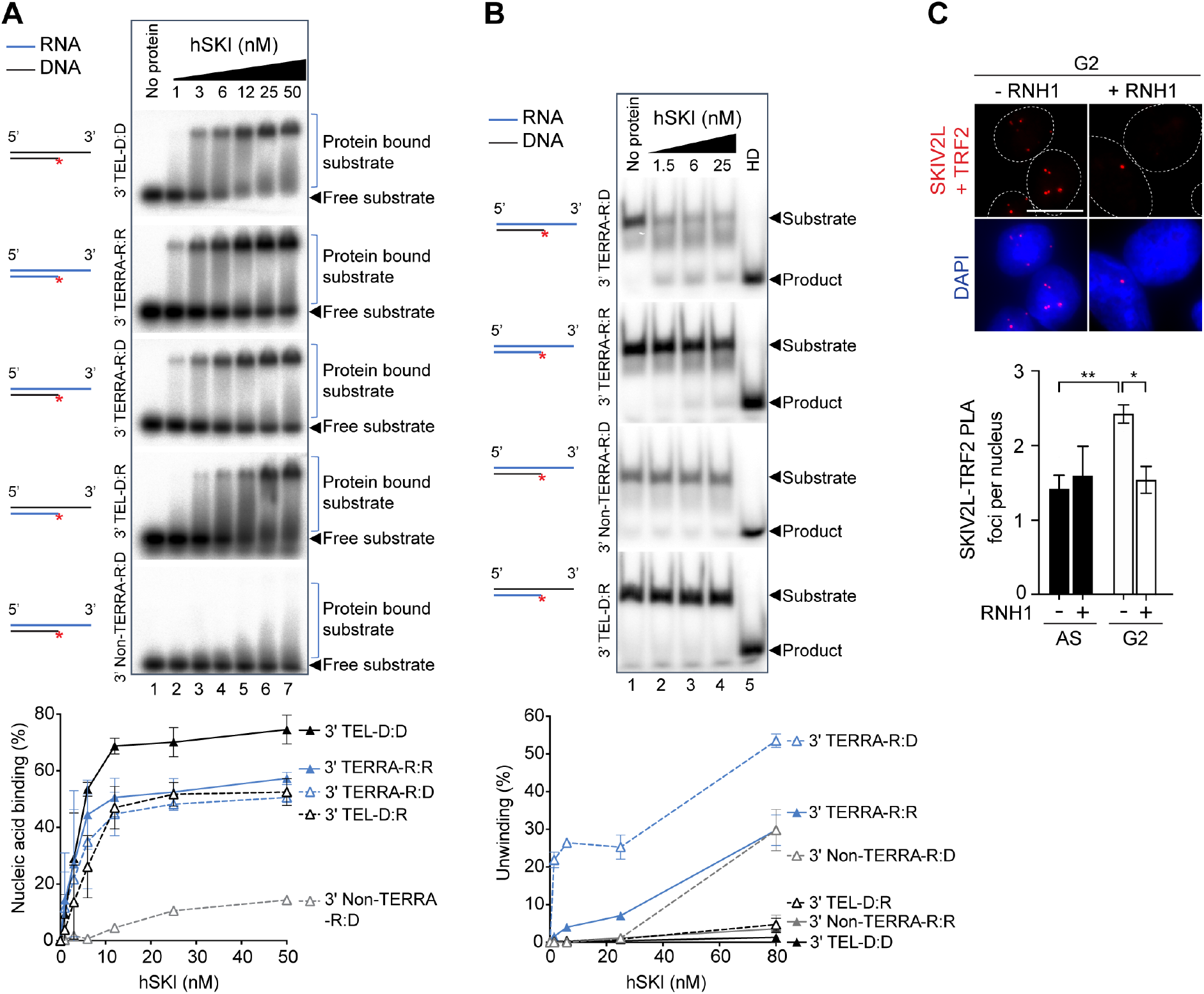
hSKI binds and unwinds telomeric DNA-RNA hybrids containing 3’ overhangs. (A) Electrophoretic mobility shift assays showing binding of hSKI to different 3’ overhang DNA/RNA (D/R) substrates and quantification (bottom) (means ± SEM, n = 2). (B) Unwinding assays demonstrating efficiency of different 3’ overhang DNA/RNA (D/R) substrates by hSKI and quantification (bottom) includes data from S4A (means ± SEM, n = 2-4). HD, heat denatured substrate. (C) Proximity ligation assay showing co-localisation of SKIV2L and TRF2 in asynchronous (AS) and in G2-synchronised HeLa cells with and without RNase H1 (RNH1) overexpression (means ± SEM, n = 3-4 independent experiments, at least 600 cells scored per condition, scale bar 10 µm).

### hSKI complex preferentially unwinds DNA-RNA hybrids with telomeric RNA overhang

The SKIV2L subunit of the complex belongs to the Ski-2 like family of RNA helicases and contains a conserved ATPase domain that is essential for the helicase function. Therefore, we confirmed that our purified hSKI complex could hydrolyse ATP and was functional *in vitro* (Fig S3B). Given that the SKI complex is conserved in evolution and yeast Ski can unwind RNA duplexes in 3’ to 5’ direction (Halbach et al. 2013), we next assessed the ability of our purified hSKI to unwind DNA-RNA duplex substrates with 3’ overhangs using *in vitro* unwinding assays. Having established the affinity of hSKI for telomeric sequences, we compared overhangs made of either RNA- or DNA-telomeric sequences (Figure 4B and S4A). Surprisingly, hSKI best unwound a DNA-RNA hybrid substrate with 3’ RNA telomeric overhang, where the substrate was unwound by low nanomolar concentration of hSKI (3’ TERRA-R:D, 20% substrate unwound with 1.5 nM protein, Figure 4B). As expected, the unwinding activity is ATP dependent (Figure S4B). Additionally, hSKI also unwound telomeric RNA duplex but with much less efficiency (3’ TERRA-R:R, 25% substrate was unwound with 80 nM protein) (Figure 4B and S4A). Similarly, DNA-RNA hybrids with non-telomeric RNA overhang were only unwound at the highest concentration of SKI (25% unwinding with 80 nM protein, Figure 4B and S4A). Other 3’ overhang substrates such as RNA duplex with non-telomeric sequence, DNA-RNA hybrid with telomeric DNA overhang and DNA duplex with telomeric overhang were all poor substrates of hSKI and could not be unwound (Figure 4B and S4A).

The preference of hSKI for unwinding DNA-RNA hybrids with telomeric RNA sequence that mimics TERRA in these assays is also validated by the binding of hSKI to these same substrates at low protein concentration (1 nM, Figure 4A). Altogether, the biochemistry indicates a binding and unwinding function of G-rich DNA-RNA hybrids by the hSKI complex and suggests a possible function *in vivo*, where it could be involved in the metabolism of such molecules at telomeres.

We next investigated whether the localisation of hSKI to telomeres is dependent on the presence of DNA-RNA hybrids. We overexpressed RNase H1-EGFP, which specifically cleaves the RNA moiety of DNA-RNA hybrids (Cerritelli and Crouch 2009) or EGFP alone as a control in HeLa cells and looked at the co-localisation of SKIV2L with TRF2 by PLA (Figure 4C and S5A). We found that RNase H1 overexpression suppresses the increase in SKIV2L-TRF2 PLA foci observed in G2-synchronised cells compared to asynchronous EGFP-transfected cells (control). This suggests that SKIV2L localisation to telomeres in G2 is dependent on telomeric DNA-RNA hybrids.

### SKIV2L facilitates telomeric DNA-RNA hybrid regulation in G2

In order to investigate the possible regulation of DNA-RNA hybrids by the hSKI complex, we used immunofluorescence of the S9.6 antibody that recognises DNA-RNA hybrids (Boguslawski et al. 1986; Garcia-Rubio et al. 2018). Notably, and in agreement with previous data (Barroso et al. 2019), HeLa cells synchronised in G2 present higher average nuclear S9.6 signal intensity than the asynchronous cell population (Figure 5A, shCtr AS *vs* shCtr G2), which remained unchanged in SKIV2L-deficient cells synchronised in G2 (Figure 5A). Surprisingly, the average nuclear intensity of DNA-RNA hybrids did not vary drastically in SKIV2L-depleted cells compared to the control cells (Figure 5A, shCtr AS *vs* shSKIV2L AS). This observation suggests that SKIV2L alone is not responsible for the regulation of genomic nuclear DNA-RNA hybrids in G2 (Figure 5A). As expected, overexpression of RNase H1 in all conditions significantly reduced the average S9.6 nuclear signal intensity.

**Figure 5.**
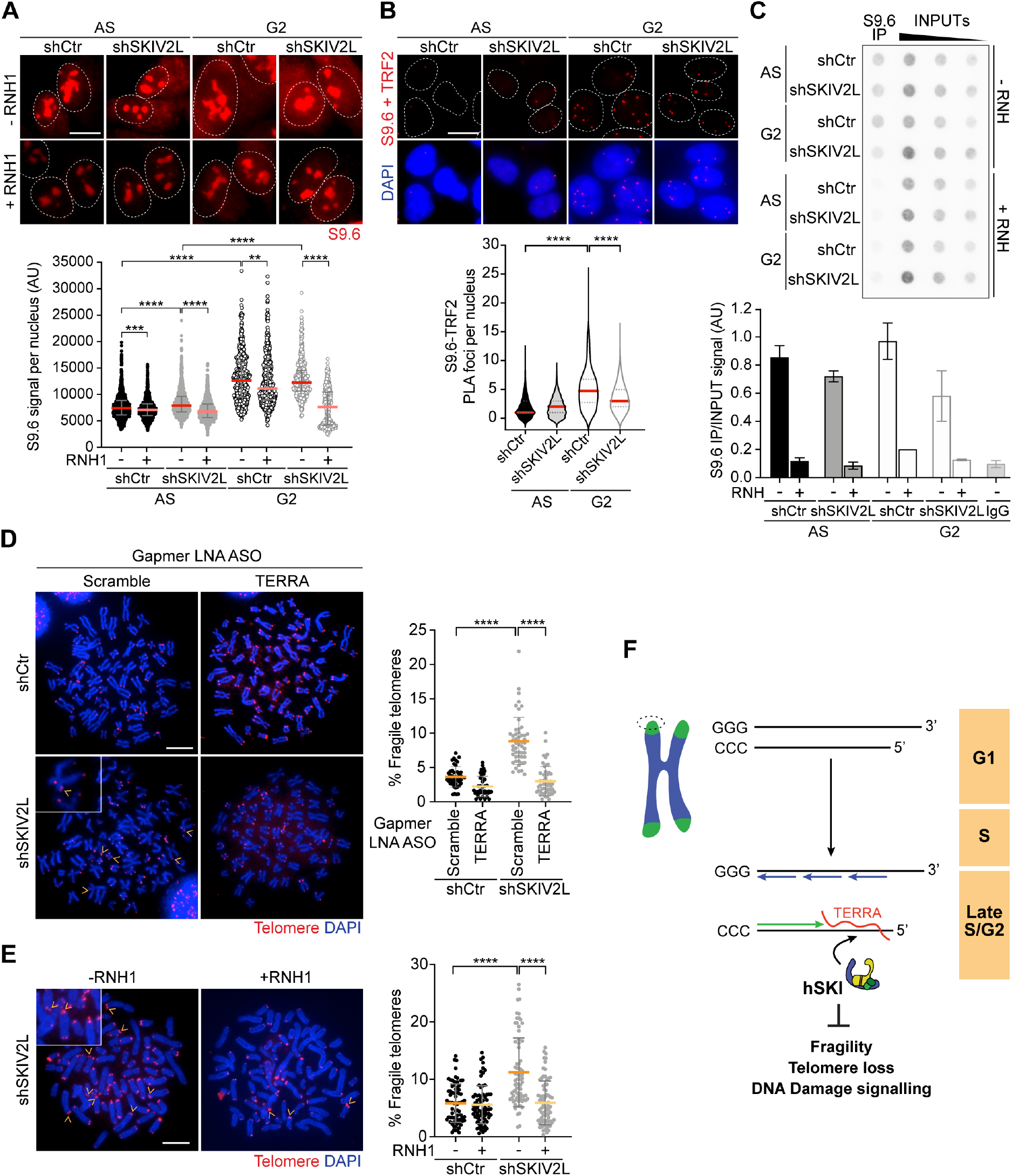
hSKI regulates telomeric DNA-RNA hybrids *in vivo* to prevent telomere fragility. (A) S9.6 IF in HeLa cells treated with RNAse III (median ± interquartile range, 2-4 independent experiments, at least 400 cells scored per condition, scale bar 10 µm). Mann-Whitney U test * p<0.05, ** p<0.01, ***p <0.001, ****p <0.0001. (B) PLA showing the co-localisation of DNA-RNA hybrids (S9.6) and TRF2 in HeLa cells pre-extracted and treated with RNAse III (median, Q1 and Q3, 2 independent experiments, at least 400 cells scored per condition, scale bar 10 µm). Mann-Whitney U test ****p <0.0001. (C) DRIP showing the amount of DNA-RNA hybrids at telomeres in AS and G2-synchronised HeLa cells, RNH, RNase H treatment (means ± SEM, n = 2 independent experiments). (D) Telomeric FISH analysis: % of telomere fragility in HeLa cells transfected with gapmer LNA targeting scramble or TERRA sequences (means ± SD, n >= 42 metaphases, 2 independent experiments, scale bar 10 µm). t test **** p <0.0001. (E) Telomeric FISH of cells with (+) or without (−) RNase H1 (RNH1) overexpression. Details are as in (D) (n >= 74 metaphases, 3 independent experiments) t test **** p <0.0001. (F) Model proposing the function of hSKI at telomeres. Telomeric DNA-RNA hybrid accumulation in late S/G2 phase drives hSKI recruitment to telomeres to regulate physiological DNA-RNA hybrid levels and ensure telomere stability.

Telomeric DNA-RNA hybrids can form by hybridisation of TERRA molecules with the C-rich telomeric leading strand (Balk et al. 2013; Pfeiffer et al. 2013). Thus, in order to assess the presence of DNA-RNA hybrids specifically at telomeres, we performed PLA using S9.6 and TRF2 antibodies and measured the resultant co-localisation signal (Figure 5B). We observed a significant decrease of PLA foci in SKIV2L-depleted cells synchronised in G2 compared to the Ctr while there was no effect in the asynchronous populations (Figure 5B). To further support this result, we carried out two independent sets of DNA-RNA immunoprecipitation experiments (DRIP, with sonicated and digested DNA), where the S9.6 antibody is used to precipitate DNA-RNA hybrids, followed by telomere dot-blot. We detected a slight increase of DNA-RNA hybrids at telomeres in G2 cells compared to control (Figure 5C), while depletion of SKIV2L led to a decrease in the amount of DNA-RNA hybrids at telomeres (Figure 5C and S5B), confirming the PLA result. Importantly, the signal was abolished in all conditions upon treatment with RNase H, confirming the specificity of the assay. RNA dot-blot and Northern blotting were used to control for telomeric RNA levels. SKIV2L depleted cells showed similar levels of global TERRAs compared to control cells (Figure S5C-D), with a slight decrease of long TERRA molecules in SKIV2L mutant (Figure S5D arrow). Collectively, our data suggest that SKIV2L is important for the protection or stabilisation of telomeric DNA-RNA hybrids and that it may facilitate their regulation at telomere chromatin.

### SKIV2L ensures telomere stability in a DNA-RNA hybrid-dependent manner

There is strong evidence that DNA-RNA hybrids are physiologically important for telomere homeostasis (Balk et al. 2013; Pfeiffer et al. 2013; Graf et al. 2017). Nevertheless, persistent or mis-regulated DNA-RNA hybrids can act as barriers to replication progression and hence, a major source of replication stress and genome instability, including telomere fragility (Toubiana and Selig 2018; Crossley et al. 2019; Garcia-Muse and Aguilera 2019; Niehrs and Luke 2020). Considering that G2 synchronised HeLa cells present SKIV2L-protected DNA-RNA hybrids necessary for telomere stability, we decided to investigate the effect of removing hybrids and TERRAs in these cells. Firstly, we downregulated TERRAs by transfecting locked nucleic acid (LNA) gapmers antisense oligonucleotides (ASOs) (Chu et al. 2017), in Ctr and SKIV2L-depleted cells. We found that telomere fragility triggered by SKIV2L knockdown was suppressed after TERRAs downregulation (Figure 5D). We next tested whether telomere fragility occurring in SKIV2L-depleted cells was dependent on the presence of telomeric DNA-RNA hybrids, by transiently overexpressing RNase H1 (Figure 5E and S5A). Consistent with the aforementioned result, overexpression of RNase H1 led to a statistically significant reduction in telomere fragility to control levels. Importantly, it suggests that telomere fragility is caused by the formation of pathological telomeric DNA-RNA hybrid structures when SKIV2L is absent. Taken together, our results implicate the hSKI complex in safeguarding telomere integrity by maintaining a proper telomere structure and preventing DNA-RNA hybrid-mediated DNA replication stress at telomeres.

## Discussion

SKI is an essential cofactor of the RNA exosome function in the cytoplasm, which ensures mRNA turnover and quality control (Wang et al. 2005; Halbach et al. 2013; Aly et al. 2016; Tuck et al. 2020; Zinoviev et al. 2020). However, and to date, no nuclear function on non-coding RNA processing has been ascribed to the hSKI complex. In this study, we describe a previously unforeseen function of hSKI in the nucleus and particularly at telomeres, where the complex localises during G2 to prevent telomere instability before mitosis. Strikingly, SKIV2L depletion in HeLa cells reduces the recruitment of TTC37 and WDR61 to telomeres, suggesting a pivotal role for SKIV2L in the stability of hSKI complex in its recruitment to telomeres.

The recruitment of hSKI to telomeres coincides with a time when TERRA levels are rising again after reaching their lowest levels in late S, in HeLa cells (Porro et al. 2010). TERRAs can pair with the telomeric C-rich DNA strand generating DNA-RNA hybrids. In addition to their physiological function to regulate telomere biology (Balk et al. 2013; Chu et al. 2017; Graf et al. 2017; Niehrs and Luke 2020), these structures may constitute obstacles for the replication fork and perturb genome stability (Crossley et al. 2019; Garcia-Muse and Aguilera 2019), particularly at telomeres (Arora et al. 2014; Sagie et al. 2017). We show that although the global levels of DNA-RNA hybrids are not overly affected, telomeric DNA-RNA hybrids levels are reduced in SKIV2L-depleted G2 cells. This suggest that the nuclear function of SKIV2L and the hSKI complex is possibly specific to telomeres and less extended to other genomic regions. Moreover, telomere fragility occurring in the absence of SKIV2L is suppressed upon TERRA downregulation or overexpression of RNase H1 ribonuclease, a factor that removes genomic and also telomeric DNA-RNA hybrids in the cells (Arora et al. 2014; Silva et al. 2019; Feretzaki et al. 2020). This indicates a key role for SKIV2L in preventing DNA-RNA hybrid-associated telomere instability.

SKIV2L recruitment to telomeres is dependent on the accumulation of DNA-RNA hybrids in G2, which is also supported by LC-MS/MS-pull-down assays with synthesised DNA-RNA hybrids, in which SKIV2L and TTC37 were identified as DNA-RNA hybrid binding factors (Wang et al. 2018). SKIV2L belongs to the superfamily 2 (SF2) of RNA helicases and has two characteristic RecA-like domains, responsible for nucleic acid binding as well as ATP hydrolysis (Fairman-Williams et al. 2010). Similar to yeast SKI, hSKI can hydrolyse ATP, an activity that is also essential for its mRNA extraction role (Halbach et al. 2013; Zinoviev et al. 2020). Our biochemical analysis shows that hSKI binds RNAs including TERRA, however the affinity is much higher for telomeric DNA, and in particular DNA-RNA hybrids with 3’ telomeric overhangs. While the overall affinity is higher for telomeric DNA, hSKI unwinds DNA-RNA hybrids containing 3’ overhang of TERRA-like RNA sequence much more efficiently than DNA overhangs. Therefore, we propose that hSKI could load itself on telomeric ssDNA and facilitate the correct processing of telomeric DNA-RNA hybrids by acting over the RNA moiety of the hybrid.

Upon its binding, hSKI might promote/facilitate the correct processing of physiological telomeric DNA-RNA hybrids possibly by preventing the action of other factors that in turn would provoke their aberrant processing into pathological intermediates, causing telomere instability. This function could be exerted by hSKI alone or in concert with other RNA regulatory factors, including 5’ to 3’ RNA helicase UPF1, hEST1A or SMG1, all involved in telomere length homeostasis by negatively regulating TERRA association with telomeres (Azzalin et al. 2007; Chawla et al. 2011).

In agreement with previous reports showing increased levels of DNA-RNA hybrids in G2 of the cell cycle (Barroso et al. 2019; Gomez-Gonzalez et al. 2020), our results also suggest that telomeres present more DNA-RNA hybrids in G2, that mediates the recruitment of additional factors, including hSKI. The physiological function of those hybrids in G2 remains to be investigated. They could be a prerequisite for mitotic telomere condensation, considering the strong correlation between DNA-RNA hybrids and chromatin compaction marks in both yeast and human cells, and their importance for the establishment of repressive heterochromatin at mammalian gene terminators (Castellano-Pozo et al. 2013; Skourti-Stathaki et al. 2014; Chedin 2016; Crossley et al. 2019). Alternatively, the equilibrium among dsDNA and DNA-RNA hybrid structures could favour the affinity of different regulatory factors to telomeres, including hSKI. Interestingly, this kind of regulation has been recently uncovered for the DNA methyltransferase 1 (DNMT1) at gene promoters (Grunseich et al. 2018). DNMT1 binding affinity to DNA-RNA hybrid structures is reduced compared to dsDNAs, thus decreasing promoter methylation and promoting transcription at R loop-enriched regions. We cannot exclude that a similar regulation, mediated by hSKI or other RNA processing factors, might occur at human telomeres so that it would affect the activity of the subset of TERRA promoters containing CpG islands, which are regulated by both DNMT1 and DNMT3b enzymes (Nergadze et al. 2009; Feretzaki et al. 2019). Interference with telomere replication is usually indicated by telomere fragility, observed as a multi-telomeric signal detected in metaphase chromosomes (Sfeir et al. 2009). The fact that telomere fragility and ATM-mediated DNA damage checkpoint activation are suppressed upon the inhibition of DNA polymerase using aphidicolin, support the conclusion that pathological intermediates resulting from an aberrant processing of DNA-RNA hybrids in the absence of hSKI do interfere with telomere DNA replication, eventually leading to DNA breaks. Notably, our results regarding DNA damage response activation upon SKIV2L depletion are consistent with published data using episomal systems to study the consequences of transcription-replication conflicts (Hamperl et al. 2017). The authors showed that upon co-directional R-loop-mediated transcription-replication conflicts, as those potentially occurring at telomeres, both ATM-Chk2 DNA damage response pathway is activated and R-loop levels are reduced, while no activation of Chk1 is observed, which is consistent with our findings.

Interestingly, the loss of telomeric signal frequently occurring in SKIV2L-depleted cells is not restored down to normal levels upon DNA replication inhibition (APH). This fact could be explained by two different scenarios: Firstly, hSKI could have different functions at telomeres depending on its recruitment by various DNA secondary structures, similar to RTEL1 (Vannier et al. 2012; Vannier et al. 2013). Secondly, hSKI-dependent regulation of telomeric DNA-RNA hybrids in G2 would ensure their physiological function in order to protect the telomere structure. Therefore, when those regulatory hybrids are not protected (*e*.*g*. SKIV2L depletion), telomere loss will still occur by inappropriate processing of aberrant DNA-RNA intermediates, which overexpression of RNase H1 would remove, hence preventing telomere fragility.

We propose a model in which hSKI has an unexpected role in safeguarding telomere stability in human cells (Figure 5F). According to our model, hSKI is recruited to telomeres in late S/G2 phase of the cell cycle in a DNA-RNA hybrid-dependent manner. Once there, hSKI facilitates the correct processing of DNA-RNA hybrid structures and prevent the accumulation of aberrant structures, which may hinder DNA replication and result in increased telomere fragility. Consequently, SKIV2L protects telomere ends from DNA damage signalling activation.

In summary, this study provides the first evidence that the human SKI complex, known for its cytoplasmic role in NMD, is also a chromatin factor. In particular, hSKI associates with telomeres and ensures telomere stability before mitosis. The regulation of telomeric lncRNAs (TERRAs) and DNA-RNA hybrids is essential for telomere homeostasis and cells use different components of the RNA processing machineries, including the hSKI complex, to achieve chromatin compaction, cell cycle regulation and genome stability. Further efforts will be necessary to fully understand the complexity and regulation of the different factors involved in telomeric RNA processing along with the human SKI complex at telomeres.

## Material and Methods

### Cell culture and generation of stable cell lines

HeLa1.3 cells referred as “HeLa” across the manuscript (kindly provided by T. de Lange), U2OS (ATCC), 293FT (ATCC) and HT1080-ST (from J. Lingner) were cultured in DMEM medium supplemented with 10% (v/v) fetal bovine serum (FBS, Sigma-Aldrich, F2442) and maintained at 37°C in 5% (v/v) CO_2_. *Spodoptera frugiperda* Sf9 insect cells (kindly provided by P. Cejka) were grown at 27°C in serum free Insect-XPRESS medium (Lonza, BELN12-730Q).

For the generation of SKIV2L and CONTROL knockdown stable cells, lentiviral particles were produced in 293FT cells using pLKO.1 SKIV2L shRNA construct (Dharmacon, 5’-GTACACTATGATCCTCAACTT-3’, RNAi Consortium TRCN0000051816) and pGIPZ (Control, Ctr) constructs. HeLa cells were transduced following standard procedures. Selection and clonal population generation was achieved with 1 µg/ml puromycin. Cells expressing the shRNA were maintained in 0.25 µg/ml puromycin-containing media.

### DNA plasmids and transfections

For the overexpression of RNase H1 we used pEGFP-M27-H1 and pEGFP as a control (kindly gifted to us from RJ. Crouch) (Cerritelli et al. 2003). Their transient transfection into HeLa cells was performed using the Nucleofector kit R (Lonza #VVCA-1001) according to the manufacturer’s instructions. Briefly, 4 million HeLa cells were transfected with 12 µg of plasmid DNA (in a final volume of 100 *μ*l in Nucleofector solution (82 µl) mixed with supplementary solution (18 µl) supplied in the kit. Nucleofector™ II/2b Device (Lonza #LO AAB-1001) programme A-020 was used. Cells were seeded into 1 × 10 cm dishes for each condition and collected after 24 h for downstream analysis.

For TERRA knockdown previously described TERRA (5′-TAACCCTAACCCTAAC-3’) and Scramble (5′-CACGTCTATACACCAC-3′) LNA gapmers were synthesised by Exiqon (Chu et al. 2017). 2 × 10^6^ HeLa cells were seeded in a 10 cm dish and transfected with 100 nM of LNA gapmers using Lipofectamine RNAiMAX (Invitrogen 13778150) in a final volume of 3 ml following the manufacturer’s instructions, incubated for 3 h before the replacement of medium and collected after 48 h.

### Cell synchronisation and drug treatments

HeLa cells were treated with 2.5 mM thymidine (Sigma-Aldrich, T1895) for 24 h, then washed and allowed to grow in normal media for 16 h followed by a second incubation with 2.5 mM thymidine for 24 h. Cells were released from the block and collected at the specified time points. S-phase or G2-phase enriched cell populations were released for 4 h and 7 h, respectively. For enrichment of mitotic cells, HeLa were initially treated with 2.5 mM thymidine for 16 h. After a release for 8 h in fresh media they were incubated with 50 ng/mL nocodazole (Sigma-Aldrich, M1404) for 16 h. Cells were collected with no release period. When indicated, HeLa cells were treated with 0.2 *μ*M aphidicolin for 20 h or with the same amount of DMSO (untreated control). If indicated, transfection of synchronized G2 HeLa cells (in parallel with asynchronous HeLa) was performed as described above in the ‘DNA plasmids and Transfection’ section after the first thymidine block, coinciding with the beginning of the first release.

### Fractionation assay

HeLa cells were scraped in ice-cold CSK buffer (10 mM PIPES KOH pH 6.8, 100 mM NaCl, 300 mM Sucrose, 1.5 mM MgCl2, 5 mM EDTA, 0.5% (v/v) Triton-X-100, cOmplete, EDTA-free Protease Inhibitor Cocktail 2X (Roche, 11873580001), phosphatase inhibitor (Sigma-Aldrich, P0044), homogenised and incubated on ice for 10 min. The 1/3 was kept as the whole cell extract. The rest of the sample was centrifuged for 3 min at 3000 rpm. The supernatant was collected as the soluble fraction, whilst the pellet was washed once with CSK buffer and resuspended in lysis buffer. This was homogenised with a 0.5 mm needle and incubated at room temperature for 20 min before snap freezing. *α*-Tubulin (cytoplasmic) and H3 (chromatin-binding) were used as controls.

### Immunofluorescence (IF)

Cells were grown on 4-well culture slides, pre-extracted for 5 min with permeabilisation buffer (50 mM NaCl, 3 mM MgCl2, 20 mM Tris pH 8, 0.5% (v/v) Triton X-100, 300 mM sucrose), fixed for 15 min in fixative solution (formaldehyde 4% (w/v) / sucrose 2% (w/v)) and permeabilised for another 10 min. Slides were incubated for 30 min with the blocking buffer (10% (v/v) goat or donkey serum (Stratech Scientific Ltd) in 1X PBS) at 37°C. Then, the primary antibody (anti-SKIV2L, Proteintech group, 11462-1-AP, 1:1000; anti-TTC37, Novus Biologicals, NBP1-93640, 1:500; anti-WDR61, Sigma-Aldrich, SAB1401852, 1:750 or anti-TRF2, Millipore, 05-521, 1:500) was added in blocking buffer and incubated for 1 hour at 37°C. After 3 x 3 min washes with 1X PBS, the secondary antibody (1/400 in blocking buffer, goat anti-rabbit Alexa 594 antibody, Thermo Fisher, A-11037 or donkey anti-mouse Alexa 488 antibody, Thermo Fisher, R37114) was added in blocking buffer for 30 min at 37°C, followed by 3 x 3 min washes with PBS 1X. Finally, mounting media containing DAPI (Prolong gold antifade mountant with DAPI, Invitrogen, P36931) was used to counterstain DNA.

### Proximity Ligation Assay (PLA)

For proximity ligation assay experiments, HeLa cells were cultured on glass coverslips and pre-extracted, fixed and permeabilised as described above. For S9.6 PLA, samples were treated with RNase III as described above. PLA was performed following the manufacturer’s instructions (Duolink DUO92101, Sigma-Aldrich) with some modifications. For S9.6-TRF2 PLA, cells were blocked for 3 h (1 h at 37°C and 2 h at 4°C) in blocking buffer (BSA 2% (w/v) in PBS), incubated with the mouse anti-DNA-RNA hybrid S9.6 antibody (1/2000, Kerafast ENH001) and rabbit anti-TRF2 antibody (1/1000, Novus NB110-57130) in blocking buffer overnight at 4°C followed by 3 washes with PBS-Tween 0.1% (v/v). For TRF2-SKIV2L PLA, IF was performed as described above using anti-SKIV2L (Proteintech group, 11462-1-AP, 1:500) and anti-TRF2 (Millipore, 05-521, 1:500) primary antibodies. For detection, anti-Mouse minus and anti-Rabbit plus PLA probes were incubated in blocking buffer for 1 h at 37°C and ligation and amplifications reactions were performed for 30 and 100 min respectively at 37°C. Slides were imaged as described above. Control reactions using only one of the antibodies in each case were performed.

### PICh and mass spectrometry analysis

PICh experiments were conducted as previously described (Dejardin and Kingston 2009; Porreca et al. 2020) with minor modifications on the sample processing. Protein samples were processed using the Filter Aided Sample Preparation (FASP) protocol (Wisniewski et al. 2009). Briefly, samples were loaded onto 30 kDa centrifugal concentrators (Millipore, MRCF0R030) and buffer exchange was carried out by centrifugation on a bench top centrifuge (15min, 12,000g). Multiple buffer exchanges were performed sequentially with UA buffer (8M urea in 100mM Tris pH 8.5, 3×200 µl), reduction with 10mM DTT in UA buffer (30min, 40°C) and alkylation with 50mM chloroacetamide in UA buffer (20min, 25°C). This was followed by buffer exchange into UA buffer (3×100 µl) and 50mM ammonium bicarbonate (3×100 µl). Digestion was carried out with mass spectrometry grade trypsin (Promega, V5280) using 1 µg protease per digest (18h, 37°C). Tryptic peptides were collected by centrifugation into a fresh collection tube (10 min, 12,000g) and washing of the concentrator with 0.5M sodium chloride (50 µl, 10 min, 12,000g) for maximal recovery. Following acidification with 1% (v/v) trifluoroacetic acid (TFA) to a final concentration of 0.2% (v/v), collected protein digests were desalted using Glygen C18 spin tips (Glygen Corp, TT2C18.96) and peptides eluted with 60% (v/v) acetonitrile, 0.1% (v/v) formic acid (FA). Eluents were then dried using vacuum centrifugation. (LC-MS/MS) analysis and raw data processing is detailed in Supplemental Materials and Methods.

### Chromatin Immunoprecipitation (ChIP)

Chromatin preparation and ChIP experiments were performed as previously described (Porreca et al. 2018) with the following modifications: Chromatin (50 µg) was diluted in 1 ml final volume of ChIP dilution buffer (20 mM Tris-HCl pH 8, 150 mM KCl, 2 mM EDTA pH 8, 1% (v/v) Triton X-100, 0.1% (w/v) SDS), pre-cleared for 2 h with Protein A Dynabeads (Invitrogen, 10001D), followed by overnight incubation with 5 µg of antibody (anti-SKIV2L Proteintech group, 11462-1-AP; anti-TRF1 SantaCruz, sc-9143). Probe preparation and membrane detection were performed as described in the RNA dot blot section (see Supplemental Materials and Methods).

### DNA-RNA Immunoprecipitation (DRIP)

DRIP experiments were performed as previously described with the following modifications (Herrera-Moyano et al. 2014; Sagie et al. 2017; Sanz and Chedin 2019). Total nucleic acids, gently extracted using phenol-chloroform from one 15-cm dish of HeLa cells growth to 75-80% confluency, were digested using BsrGI, EcoRI, HindIII, SspI and XhoI restriction enzymes (New England BioLabs, 44 U each). Digested nucleic acids were EtOH-precipitated with 1.5 μl of glycogen (Thermo Fisher Scientific, R0561) and gentle resuspended in 50 μl of 1x TE buffer. 8 μg of digested nucleic acids were treated or not with 10 μl of RNase H (New England BioLabs, M029L) overnight at 37 °C in 1x RNAse H buffer and 1/10 of the samples were used as input. Samples were incubated with 20 μl of the S9.6 antibody (Kerafast ENH001) for 14-17 h at 4°C followed by the incubation with 100 μl of Dynabeads Protein A + G (2:1 A:G proportion, Invitrogen, 10001D and 10004D) for 2 h at 4°C. Then, precipitated samples were eluted in 300 μl of elution buffer (50 mM Tris, pH 8.0, 10 mM EDTA, pH 8.0, 0.5% (v/v) SDS), treated with 5.8 μl of proteinase K (24 mg/mL, P4850) for 45 min at 55°C followed by 50 μg/ml RNase A (Roche, 10109169001) for 1 h at 37°C plus 1 h at 65°C. Finally, cleaned samples were resuspended in 100 μl 1X TE buffer and blotted using a dot-blot apparatus. Probe preparation and membrane detection were performed as described in the RNA dot blot section (see Supplemental Materials and Methods). Telomeric G-rich probe ATT(GGGATT)_4_ was used for the detection of the precipitated DNA.

### Preparation of SKI expression constructs

The *SKIV2L, TTC37* and *WDR61* genes were synthesized and cloned into pFastbac1 vectors by GenScript with the addition of 6x histidine tag and 3x flag tag before the *TTC37* gene creating pFB-SKIV2L, pFB-His-Flag-TTC37 and pFB-WDR61 plasmids respectively. All three genes were codon-optimized for expression in insect cells. The pFB-SKIV2L plasmid was further modified by adding a maltose binding protein tag (MBP) before SKIV2L gene making pFB-MBP-SKIV2L plasmid. The plasmids contained a PreScission protease cleavage site in between the tags and the gene. The cloned genes were expressed using bac-to-bac baculovirus expression system in *Spodoptera frugiperda* Sf9 insect cells. Baculoviruses were produced and the cells were infected with optimal ratios of all three viruses and incubated for 52 h. The cells were collected, washed with phosphate buffered saline and frozen in −80°C until purification.

### Purification of SKI complex

The frozen cell pellet was resuspended in 3 volumes of lysis buffer containing 50 mM Hepes-KOH pH 7.6, 1 mM DTT, 1 mM EDTA, 1:400 protease inhibitor cocktail (P8340, SIGMA), 1 mM PMSF and 30 *μ*g/ml Leupeptine for 20 mins at 4°C with gentle agitation. Then, glycerol (final concentration 16% v/v) and NaCl (final concentration 325 mM) were added and the suspension was stirred slowly for further 30 min. The suspension was centrifuged at 48,000g for 30 min to obtain the soluble extract. Next, the clarified extract was incubated with pre-equilibrated amylose resin (NEB) for 60 mins. The protein bound amylose resin was washed with MBP wash buffer containing 50 mM Hepes-KOH pH 7.6, 2 mM Beta-mercaptoethanol, 250 mM NaCl, 10 % v/v glycerol, 1 mM PMSF and 10 µg/ml Leupeptine followed by elution with MBP wash buffer supplemented with 10 mM Maltose. The MBP eluate was then incubated with pre-equilibrated nickel-nitriloacetic acid resin (Ni-NTA agarose, Qiagen) for 60 mins in the presence of 20 mM imidazole. The Ni-NTA resin was washed with Ni-NTA wash buffer containing 50 mM Hepes-KOH pH 7.6, 2 mM Beta-mercaptoethanol, 150 mM NaCl, 10 % v/v glycerol, 0.5 mM PMSF, 40 mM imidazole and eluted with Ni-NTA wash buffer containing 400 mM imidazole. The Ni-NTA eluate was then incubated overnight with PreScission protease to cleave the tags. The cleaved eluate was concentrated using a centrifugal filter (50 kDa cut-off, Amicon) and then loaded onto a Superose 6 Increase 10/300 GL column (GE Healthcare). The peak fraction containing recombinant SKI complex was collected, sub-aliquoted, snap frozen and stored at −80°C until use.

### Nucleic acid binding assay

The Electrophoretic mobility assay is carried out in a total reaction of 15 µl in a buffer containing 25 mM Hepes-KOH pH 7.6, 2 mM magnesium acetate, 50 mM NaCl, 1 mM DTT, 0.1 mg/ml BSA, 1 nM DNA/RNA substrate (in molecules) and indicated amounts of SKI protein complex. The reactions were assembled on ice, incubated at 37°C for 30 mins and products were separated on a 0.7% w/v agarose gel at 100 V for 60 mins at 4°C. The gel was dried on Hybond-XL paper (GE Healthcare) in gel dryer (Biorad), exposed to phosphor screen (GE Healthcare) and scanned on FLA-5000 (Fujifilm). The gels were quantitated using Imagequant software (GE Healthcare) and fraction of protein bound DNA for each substrate was plotted.

### Statistical analysis

Statistical analysis was performed using GraphPad Prism. Error bars, statistical methods and n, are described in figure legends. Figures were prepared using Adobe Illustrator.

## Acknowledgements

We thank Titia de Lange for providing HeLa1.3 cells, Petr Cejka for Sf9 insect cells, Joachim Lingner for HT1080-ST cells and Robert J Crouch for RNase H1-overexpressing plasmids. Special thank you to Jesus Gil, Simon Boulton and Andrés Aguilera for constructive discussions.

Vannier lab’s work is supported by the London Institute of Medical Sciences (LMS), which receives its core funding from UKRI (MRC) and by an ERC Starter Grant (637798; MetDNASecStr). Roser Gonzalez-Franco received an MRC funded PhD fellowship, Jean-Baptiste Vannier and Lepakshi Ranjha are funded by UKRI (MRC). Emilia Herrera-Moyano, Rosa Maria Porreca, Eleni Skourti are funded by ERC Starter Grant (637798; MetDNASecStr). Manos Stylianakis is funded by an MRC PhD fellowship.

## Author contributions

E.H.M, R.M.P, L.R, R.G.F, E.S and J.B.V designed the project and wrote the manuscript. Molecular biology experiments were carried out by E.H.M, R.M.P, R.G.F, E.S and M.S. Biochemistry was conducted by L.R and Y.S. A.M and H.K performed mass spectrometry running and primary analysis.

