## Supplementa Material for "The human SKI complex prevents DNA-RNA hybrid-associated telomere instability"

**Inventory of Supplemental Material:**

Supplemental Figures S1-S5

Supplemental Tables S1-S2

Table S1. DNA/RNA substrates used in this study

Table S2. Oligonucleotides used in this study

Supplemental Materials and Methods

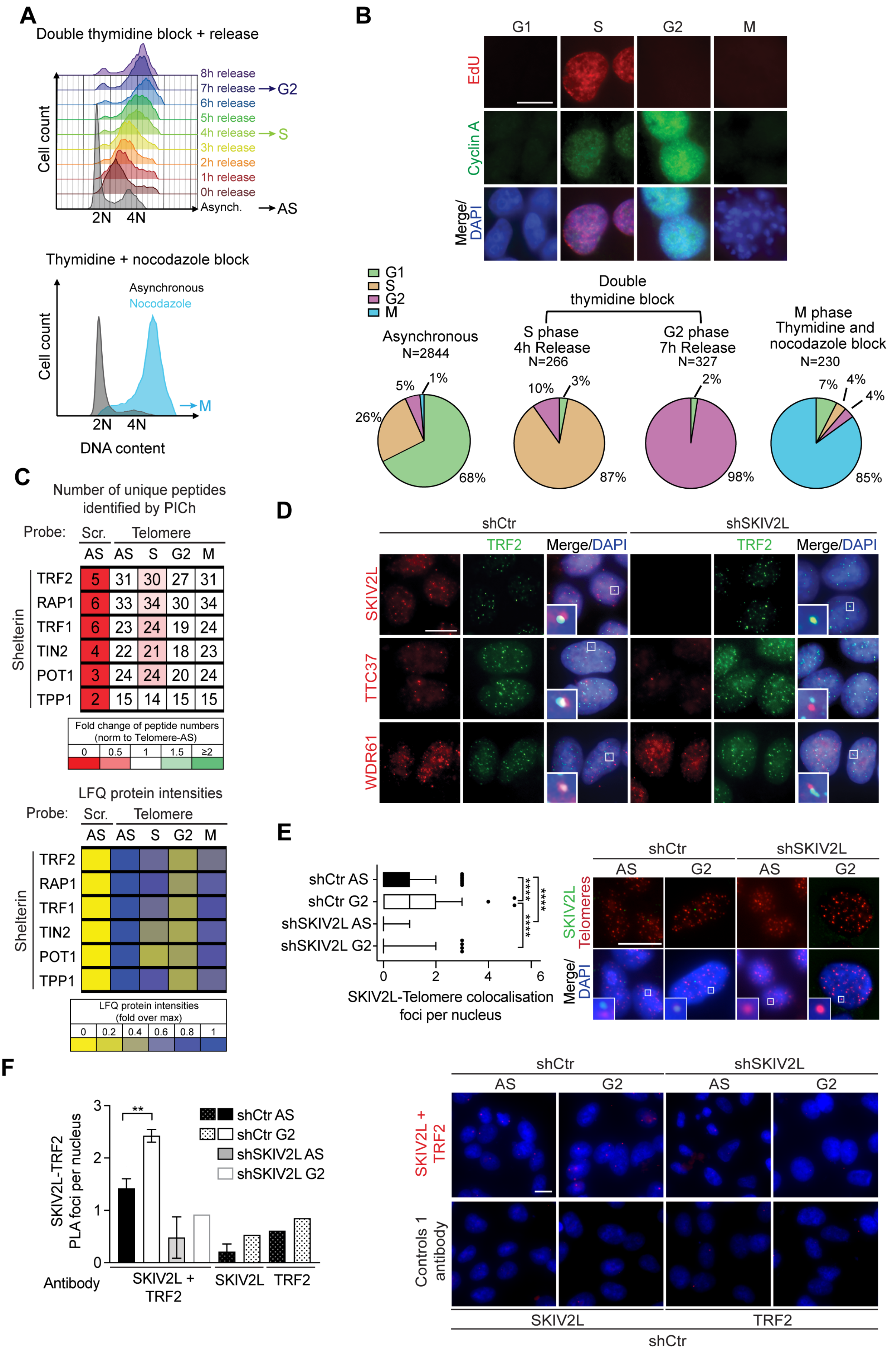

A
Double thymidine block + release
Cell count
8h release
7h release → G2
6h release
5h release
4h release → S
3h release
2h release
1h release
0h release
Asynch. → AS
2N 4N
Thymidine + nocodazole block
Cell count
Asynchronous
Nocodazole
→ M
2N 4N
DNA content
B
G1 S G2 M
EdU
Cyclin A
Merge/DAPI
G1
S
G2
M
Double thymidine block
Asynchronous N=2844
S phase 4h Release N=266
G2 phase 7h Release N=327
M phase Thymidine and nocodazole block N=230
5% 1% 26% 68%
10% 3% 87%
2% 98%
7% 4% 4% 85%
C
Number of unique peptides identified by PICh
Probe: Scr. Telomere
AS AS S G2 M
Shelterin
TRF2 5 31 30 27 31
RAP1 6 33 34 30 34
TRF1 6 23 24 19 24
TIN2 4 22 21 18 23
POT1 3 24 24 20 24
TPP1 2 15 14 15 15
Fold change of peptide numbers (norm to Telomere-AS)
0 0.5 1 1.5 ≥2
LFQ protein intensities
Probe: Scr. Telomere
AS AS S G2 M
Shelterin
TRF2
RAP1
TRF1
TIN2
POT1
TPP1
LFQ protein intensities (fold over max)
0 0.2 0.4 0.6 0.8 1
D
shCtr
shSKIV2L
TRF2
Merge/DAPI
SKIV2L
TTC37
WDR61
E
shCtr AS
shCtr G2
shSKIV2L AS
shSKIV2L G2
****
****
****
0 2 4 6
SKIV2L-Telomere colocalisation foci per nucleus
shCtr
shSKIV2L
AS G2 AS G2
SKIV2L Telomeres
Merge/DAPI
F
SKIV2L-TRF2 PLA foci per nucleus
0 1 2 3
**
shCtr AS
shCtr G2
shSKIV2L AS
shSKIV2L G2
Antibody
SKIV2L + TRF2
SKIV2L
TRF2
shCtr
shSKIV2L
AS G2 AS G2
SKIV2L + TRF2
Controls 1 antibody
SKIV2L
TRF2
shCtr

**Figure S1. Synchronisation of HeLa cells and recruitment of hSKI to telomeres at different stages of the cell cycle.** (A) FACS analysis of asynchronous (AS, black) or synchronised (coloured) HeLa cells using double thymidine block. Cells were released from the block and collected every hour (for 8 h) for FACS as specified (top panel). FACS analysis of asynchronous (black) and nocodazole blocked (M, blue) HeLa cells after an 8 h release from thymidine arrest (bottom panel). (B) Representative images of Cyclin A immunofluorescence (S-G2 phases) and EdU click-it reaction (S phase) in asynchronous HeLa cells upon incubation with EdU for 1h (top panel). Discrimination between G1 and M phases was based on DAPI signal. Scale bar, 15 µm. Quantification of IF images from AS, synchronised in S phase (4 h release), in G2 (7 h release) and in prometaphase (after 16 h nocodazole block) HeLa cells (bottom panel). Total number of quantified nuclei per condition (N) and the percentage of cells in each cell cycle phase is indicated. (C) Proteomics of isolated chromatin segments (PICh) analysis showing the binding of Shelterin proteins throughout the cell cycle (AS; S; G2; M). Tables are listing the number of unique peptides isolated including the fold change values of unique peptides normalised to the asynchronous values (top panel) and the relative LFQ intensity values identified by PICh (bottom panel). (D) Representative images of immunofluorescence showing co-localisation of hSKI with TRF2 (telomeres) in asynchronous control (shCtr) and SKIV2L-depleted (shSKIV2L) cells. Details of colocalising or non-colocalising signals are outlined. (E) IF-FISH showing co-localisation of SKIV2L and telomeres in asynchronous and G2-synchronised control (shCtr) and SKIV2L-depleted (shSKIV2L) cells (median, Q1 and Q3, n >= 210 cells, scale bar 15 µm). Mann-Whitney U test ****p <0.0001. (F) Controls for Proximity ligation assay experiment of SKIV2L-TRF2 in Figure 1E. Negative controls of PLA foci number per nucleus in shSKIV2L cells and PLA reactions using only one of the two antibodies are depicted (means ± SEM, n = 1-4 independent experiments).

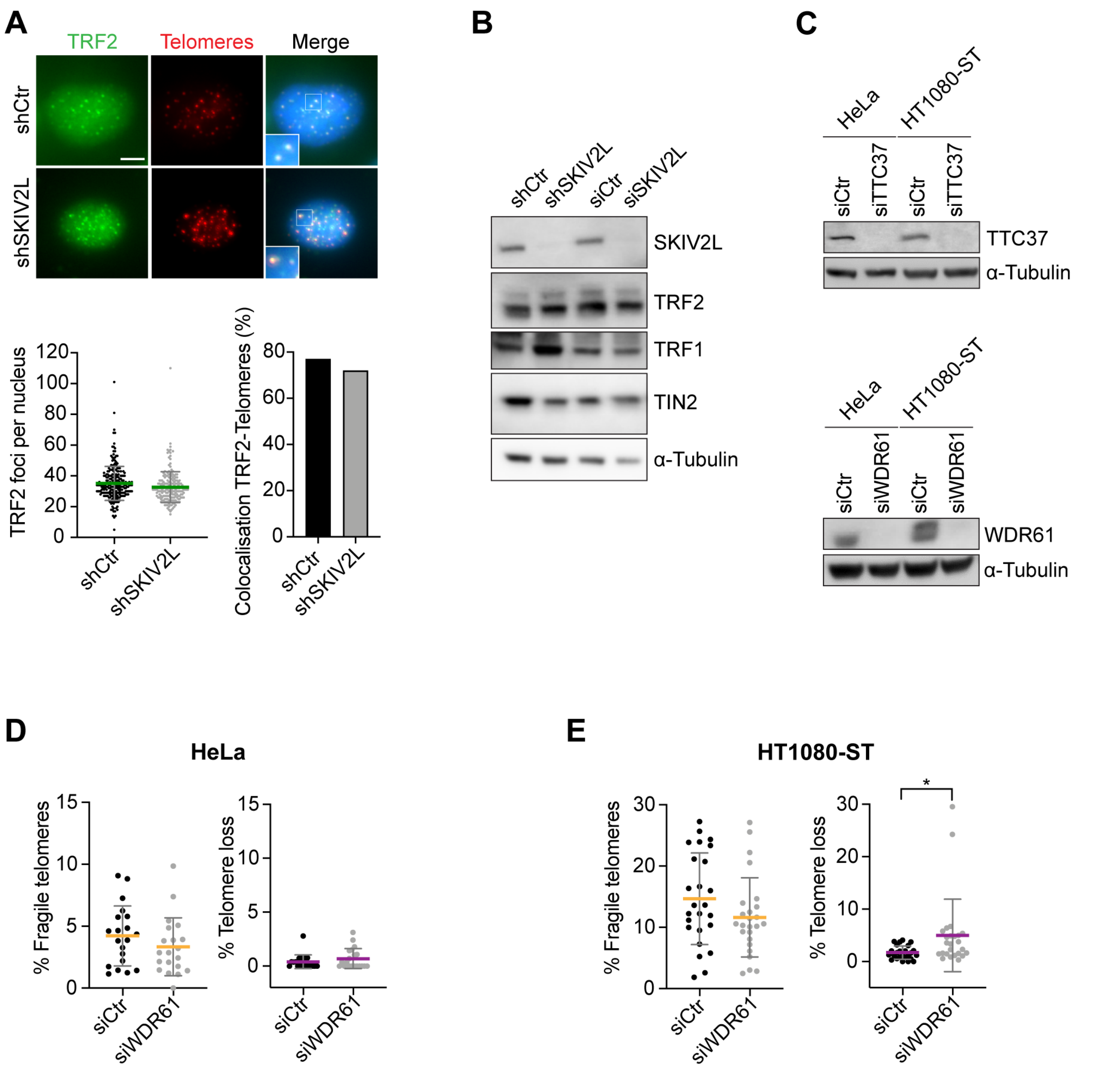


**Figure S2. Depletion of SKIV2L does not perturb Shelterin protein levels nor their binding to telomeres.** (A) IF-FISH showing number of TRF2 foci per nucleus (mean ± SD) and % of TRF2-telomeres co-localisation in control (shCtr) or SKIV2L-depleted (shSKIV2L) cells (n = 200 cells, scale bar 5 µm). (B) WB showing protein levels of SKIV2L, TRF2, TRF1 and TIN2 in control cells (shCtr/siCtr) or depleted for SKIV2L (shSKIV2L/siSKIV2L). α-Tubulin was used as a loading control. (C) WB showing protein levels of TTC37 and WDR61 in HeLa and HT1080-ST cells upon deletion of TTC37 or WDR61. α-Tubulin was used as a loading control. (D-E) Telomere FISH analysis in HeLa and HT1080-ST cells using siCtr or siWDR61: % of telomere fragility (yellow) and loss (purple) per metaphase are represented (means ± SD, n = 20-25 metaphases). t test * $p < 0.05$.

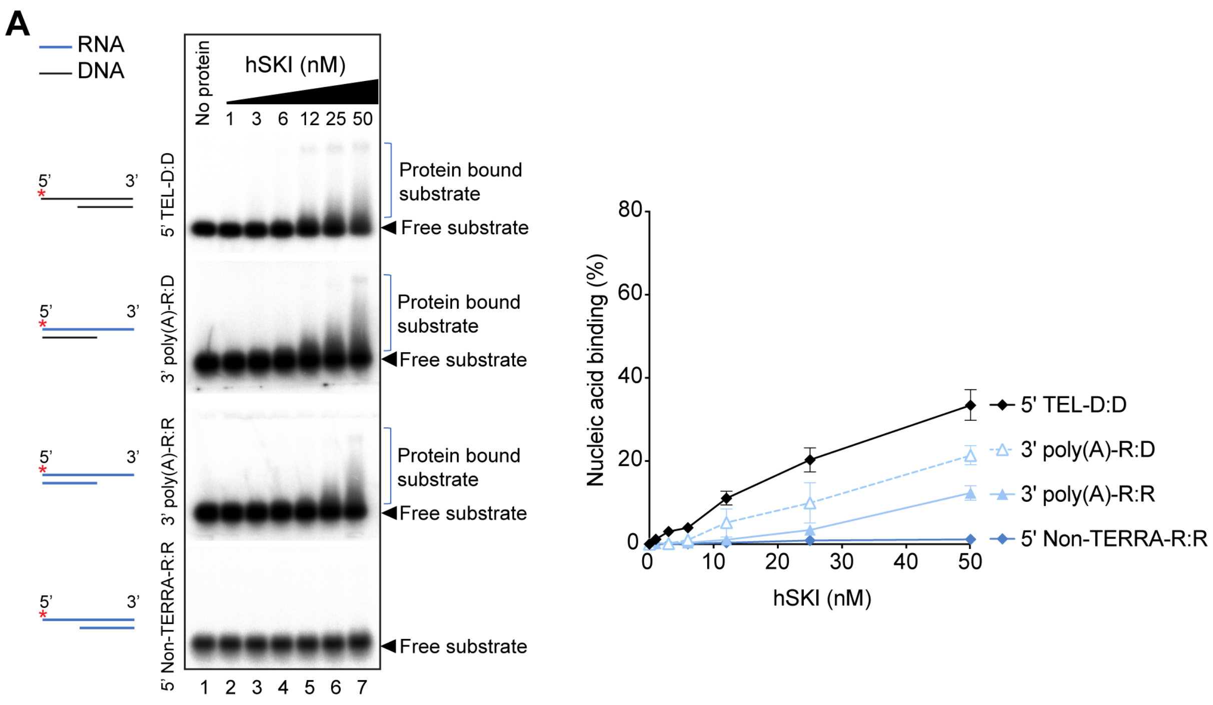


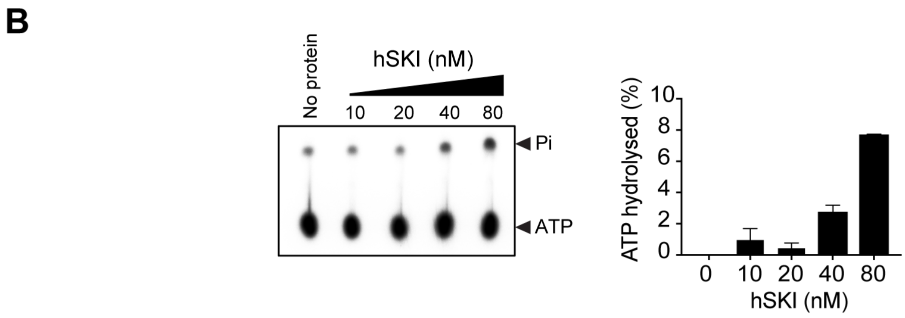


**Figure S3. Purified hSKI complex hydrolyses ATP and preferentially binds 3' telomeric overhangs.** (A) (Left) Electrophoretic mobility shift assays showing binding of hSKI complex to different 3' and 5' overhang RNA/DNA substrates. Asterisk indicates radioactive $^{32}$P label. hSKI protein amounts are as indicated (nM). (Right) Quantification of experiments in left showing % of nucleic acid binding calculated as protein bound substrate signal relative to free substrate signal (mean ± SEM, n = 2). (B) ATP hydrolysis assay showing ATPase activity of hSKI complex. The % of ATP hydrolysed is calculated as the amount of hydrolysed phosphate released (Pi) relative to ATP (mean ± SEM, n = 2).

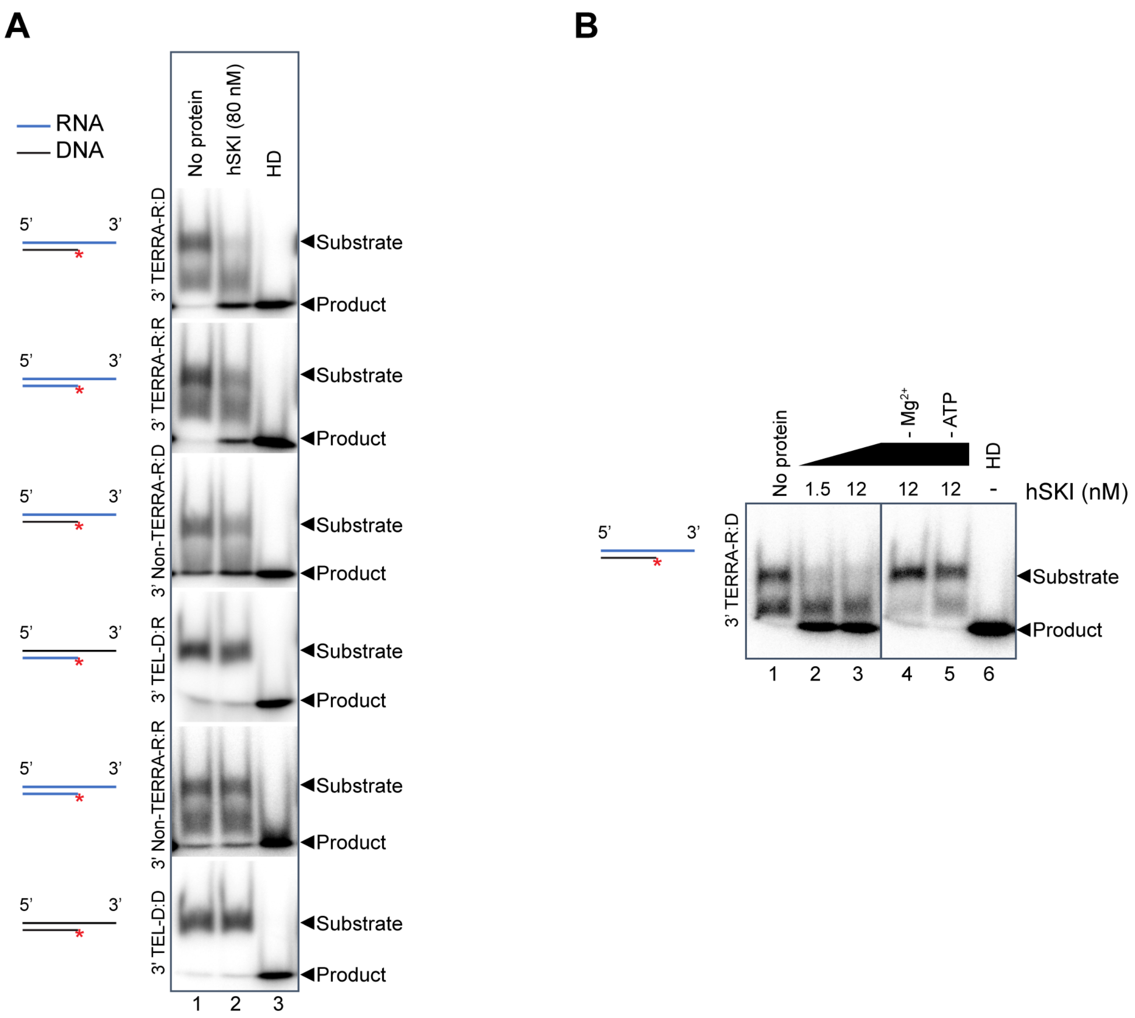


**Figure S4. Purified hSKI complex unwinds 3' TERRA overhangs.** (A) Unwinding assay of hSKI (80 nM) with various DNA/RNA substrates. HD, heat denatured substrate. (B) Unwinding assay of hSKI with 3' TERRA-RNA-DNA hybrid substrate (0.1 nM substrate, in molecules, was used here) in the absence of either magnesium (- $Mg^{2+}$) or ATP (- ATP). HD, heat denatured substrate.

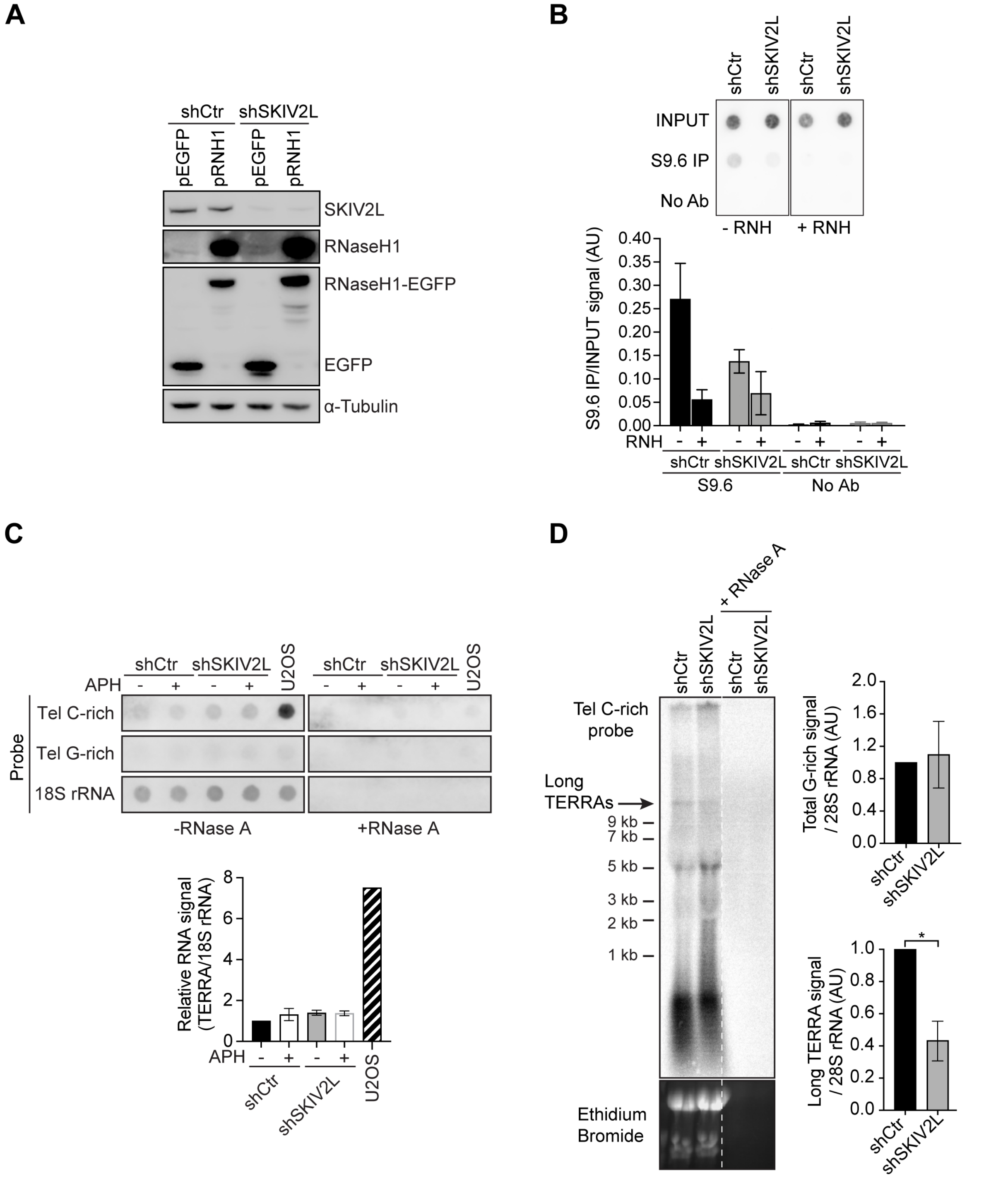


**Figure S5. Depletion of SKIV2L leads to a decrease in long TERRA levels without affecting total TERRA amounts.** (A) WB analysis showing the levels of RNase H1-EGFP (using anti-GFP and anti-RNase H1 antibodies), EGFP (using anti-GFP antibody) and SKIV2L (using anti-SKIV2L antibody) upon transfection of shCtr and shSKIV2L HeLa cells with the pRNH1 or pEGFP constructs. α-tubulin was used as a loading control. (B) DRIP showing the amount of DNA-RNA hybrids at telomeres in AS HeLa cells using sonicated nucleic acids. RNH, RNase H treatment (means ± SEM, n = 4 independent experiments). Immunoprecipitations without using antibodies were using as a negative control (n = 2). (C) RNA dot blot analysis in shCtr and shSKIV2L HeLa cells untreated or treated with aphidicolin (APH). Half of each sample was treated with Ribonuclease A (+ RNase A) before spotting to control for DNA contamination. The blot was revealed with DIG-Telomeric C-rich, Telomeric G-rich (negative control) or 18S rRNA (loading control) probes. TERRA signals were normalised to 18S rRNA signals, relative to shCtr untreated (mean ± SEM, n = 2 independent experiments). RNA from U2OS cells was used as a positive control. (D) TERRA detection by Northern blotting in shCtr and shSKIV2L HeLa cells. DIG-Tel C-rich probe was used to detect TERRA. Ethidium bromide staining of the 28S rRNA was used as a loading control. Arrow is pointing to the long TERRA band. Half of each sample was treated with Ribonuclease A (+ RNase A) to control for DNA contamination. Total TERRA signals (up) or long TERRA RNA signals were normalised to 28S rRNA signals, relative to shCtr untreated (mean ± SEM, n = 1-3 independent experiments).

**Table S1. DNA/RNA substrates used in this study**

| | Duplex substrates | Oligonucleotides used |
|---|---|---|
| 1 | dsRNA | ss poly(A) + Css27 |
| 2 | dsDNA | TEL30 + CTEL30 |
| 3 | 3’ TEL- D:D | cLR01_20 + LR01 |
| 4 | 3’ TERRA- R:R | cLR01_20 RNA + LR01 RNA |
| 5 | 3’ TERRA- R:D | cLR01_20 + LR01 RNA |
| 6 | 3’ TEL- D:R | cLR01_20 RNA + LR01 |
| 7 | 3’ Non-TERRA- R:D | 5’ss27 + Css17 DNA |
| 8 | 3’ poly(A)- R:D | ss poly(A) + Css17 DNA |
| 9 | 3’ poly(A)- R:R | ss poly(A) + Css17 |
| 10 | 3’ Non-TERRA- R:R | 5’ss27 + Css17 |
| 11 | 5’ TEL- D:D | TEL30 + 5’CTEL20 |
| 12 | 5’ Non-TERRA- R:R | 5’ss27 + 5’Css17 |

**Table S2. Oligonucleotides used in this study**

| | | Name | Sequence 5’ to 3’ |
|---|---|---|---|
| 1 | DNA | TEL18 | GGTTAGGGTTAGGGTTAG |
| 2 | DNA | Non-TEL18 | AGCGTATCCGTTCAGTTG |
| 3 | DNA | TEL30 | GGTTAGGGTTAGGGTTAGGGTTAGGGTTAG |
| 4 | DNA | CTEL30 | CTAACCCTAACCCTAACCCTAACCCTAACC |
| 5 | DNA | 5’CTEL20 | CTAACCCTAACCCTAACCCT |
| 6 | DNA | ss poly(A) DNA | CCCCACCACCATCACTTAAAAAAAAAA |
| 7 | DNA | Css17 DNA | AAGTGATGGTGGTGGGG |
| 8 | DNA | LR01 | ACGTCAGATGATCGATGGGGTTAGGGTTAG |
| 9 | DNA | cLR01_20 | CCCCATCGATCATCTGACGT |
| 10 | RNA | ss poly(A) | CCCCACCACCAUCACUUAAAAAAAAAA |
| 11 | RNA | Css27 | UUUUUUUUUUAAGUGAUGGUGGUGGGG |
| 12 | RNA | Css17 | AAGUGAUGGUGGUGGGG |
| 13 | RNA | 5’ss27 | CCCCACCACCAUCACUUACCAUCACUU |
| 14 | RNA | 5’Css17 | AAGUGAUGGUAAGUGAU |
| 15 | RNA | TERRA | UUAGGGUUAGGGUUAGGGUUAGGGUUAGGG |
| 16 | RNA | LR01 RNA | ACGUCAGAUGAUCGAUGGGGUUAGGGUUG |
| 17 | RNA | cLR01_20 RNA | CCCCAUCGAUCAUCUGACGU |

### SUPPLEMENTAL MATERIALS AND METHODS

#### FACS analysis

HeLa cells were incubated with 10 µM of EdU (Thermo Fisher, E10415) for 1 h prior to collection. Then, cell suspensions were fixed with formaldehyde 4% (w/v) in PBS for 15 min with vigorous vortexing to dissociate any remaining cellular clumps; for the same purpose the cells were passed through a syringe with a 0.5 mm needle tip prior to fixation. The Click-iT EdU Alexa Fluor 647 Flow Cytometry Assay kit (Thermo Fisher, C10634) was then used according to the manufacturer's instructions. We counterstained DNA with 10 μg/ml of Propidium Iodide (PI) (Sigma-Aldrich, P4864). Cell preparations were analysed with a BD LSR II flow cytometer.

#### Cell cycle study

Cells were seeded at a 40% confluence in 4-well slides. After the appropriate synchronisation protocol, cells were incubated for 1 h with 10 µM of EdU before collection. No pre-extraction was performed. Furthermore, for collection of mitotic enriched populations, cells were fixed with 2X fixative solution in media for 15 min to prevent excessive detachment of cells. Afterwards, the Click-iT Edu Alexa Fluor 647 Imaging Kit (Thermo Fisher, C10640) was employed to label DNA synthesis following the manufacturer's indications. Then, a regular immunofluorescence using a primary mouse anti-Cyclin A antibody (1/50, Santa Cruz biotechnology, sc-53227) was performed.

#### siRNA-mediated depletion

For the siRNA transfections, 1 million HeLa cells or 400 thousand HT-1080-ST cells were seeded per 10 cm dish 5 h before transfection. Cells were transfected with 15 μl of 20 μM siRNA (SMARTpool: ON-TARGETplus SKIV2L siRNA (Dharmacon #L-013435-01-0010), Set of 4: ON-TARGETplus TTC37 siRNA (Dharmacon #L-020959-02-0020), Set of 4: ON-TARGETplus WDR61 siRNA (Dharmacon #LQ-014614-01-0002) or ON-TARGETplus Non-targeting Pool (Dharmacon #D-001810-10-20)) using 30 μl of Lipofectamine RNAiMax transfection agent (ThermoScientific #13778150) according to manufacturer's instructions in a final volume of 3 ml (final siRNA concentration: 100 nM). Transfection was repeated 72 h later. 72 h after the second transfection, cells were harvested for downstream analysis at this time point.

**Western blotting (WB)**

Cells were trypsinised, washed in PBS 1X and spun down at 300 g for 5 min. Cell pellets were subjected to lysis by incubating in lysis buffer (NaCl 40 mM, Tris 25 mM, pH 8; $MgCl_2$ 2 mM; SDS 0.05% (w/v), Benzonase 100 units/ml (Sigma-Aldrich, E1014-25KU) and cOmplete, EDTA-free Protease Inhibitor Cocktail 2X) for 10 min on ice. Phosphatase inhibitors were added when analysing a phosphorylated protein. The lysates were sheared by being forced through a 25G needle for 10 times and then incubated on ice for 10 min. Protein concentration was determined using the Bio-Rad Protein Assay Dye Reagent Concentrate (Biorad, 500-0006) according to the manufacturer's instructions. Protein lysates were denatured for 5 min at 100ºC after addition of Laemmli buffer 4X (50 mM Tris pH 6.8; 100 mM DTT; 2% (w/v) SDS; 0.1% (w/v) bromophenol blue; 10% glycerol (v/v)), separated on NuPAGE 4-12% (w/v) Bis-Tris gels and transferred onto a nitrocellulose membrane (Amersham Protran 0.2 μm, GE10600001). The membrane was blocked using non-fat milk 5% (w/v) in PBS-T (PBS 1X; 0.1% (v/v) Tween-20) and subsequently incubated overnight at 4ºC with the primary antibody (anti-RNAseH1 (1/500, Abnova, H00246243-M01), anti-GFP (1/5000, Abcam, ab290), anti-β-actin (1/5000, Abcam, ab8226, used as a loading control), anti-ATR (1/500, Bethyl A300-138A), anti-CHK1 (1/500, Sigma-Aldrich, C9358), anti-CHK1 pS345 (1/500, Cell signaling, 2348), anti-ATM (1/500, Sigma-Aldrich, A1106), anti-ATM pS1981 (1/500, Cell signaling 4526), CHK2 (1/500, Millipore, 05-649), anti-CHK2 pT68 (1/500, Cell signaling, 2661), anti-H3 (1/1000, Abcam, ab10799), anti-SKIV2L (1/1000, Proteintech group, 11462-1-AP), anti-TTC37 (1/500, Novus Biologicals, NBP1-93640), anti-WDR61 (1/500, Sigma-Aldrich, SAB1401852; 1/1000, Thermofisher PA5-40079), anti-α-Tubulin (1/1000, Sigma-Aldrich, T6199), anti-TRF2 (1/2000, Novus Biologicals, NB110-57130), anti-RAP1 (1/500, Bethyl, A300-306A), anti-TRF1 (1/1000, SantaCruz, sc-9143), anti-TIN2 (1/1000, Sigma-Aldrich, SAB4200108)) diluted in non-fat milk or BSA 5% (w/v) in PBS-T. Subsequently, the membrane was washed 3 x 10 min in PBS-T. Following incubations with HRP-conjugated secondary antibody (anti-mouse and anti-rabbit, Dako, P0447 and P0217) in non-fat milk 5% (w/v) in PBS-T and 3 x 10 min washes in

PBS-T, the signal was visualized using ECL Western blotting reagents (Sigma-Aldrich, RPN2106) and either X-ray film exposure (Amersham Hyperfilm ECL Sigma-Aldrich, GE28-9068-35) or Amersham Imager 680 (GE Healthcare).

**IF-FISH**

For IF-FISH experiments, slides were post-fixed using fixative solution (formaldehyde 4% (w/v) / sucrose 2% (w/v)) after secondary antibody and PBS washes treatments. FISH was then performed as in metaphases, using a Cy3-O-O-$(CCCTAA)_3$ probe (PNA bio).

**Quantitative-FISH analysis (Q-FISH)**

For metaphase preparation, HeLa or HT1080-ST cells were incubated for 1 h with colcemid 10 ng/ml (Roche, 10295892001). Subsequently, cells were collected and incubated at 37°C in hypotonic buffer (HeLa: 15 min, KCl 75 mM: NaCitrate 8g/L: ddH2O, 1:1:1 and HT1080-ST: 40 min, NaCitrate 8 g/L). Fixation was performed with freshly prepared fixative (ethanol: glacial acetic acid (3:1)) followed by three washes using the same fixative. Finally, metaphase suspensions were dropped on previously humidified glass slides. Q-FISH was performed as previously described (Ourliac-Garnier and Londono-Vallejo 2011). Briefly, metaphase spreads were fixed in formaldehyde 4% (w/v) for 2 min, washed 3 x 5 min in PBS 1X, treated with pepsin (1 mg/ml in 0.05 M citric acid pH 2) for 10 min at 37°C, post-fixed for 2 min, washed and incubated in increasing ethanol concentration baths. Each slide was then covered with hybridising solution containing Cy3-O-O-(CCCTAA)3 probe (PNA bio) in formamide 70% (v/v), 10 mM Tris pH 7.4 and 1% (v/v) blocking reagent (Roche, 11096176001). This step was followed by denaturation for 3 min at 80°C on a heat block. Hybridisation was performed for 2 h at room temperature. Slides were washed 2 x 15 min in formamide 70% (v/v) in 20 mM Tris pH 7.4, followed by 3 x 5 min washes in 50 mM Tris pH 7.4, 150 mM NaCl, Tween-20 0.05% (v/v), dehydrated in successive ethanol baths and air-dried. Slides were mounted in antifade reagent containing DAPI and imaged as described above. Telomeric signals were quantified using the ImageJ software (Fiji).

**S9.6 IF**

S9.6 IF was performed as previously described (Garcia-Rubio et al. 2018; Barroso et al. 2019) with minor modifications. Briefly, cells were pre-extracted for 3 min using ice-cold pre-extraction buffer (0.5% Triton X-100, 20 mM HEPES-KOH (pH7.9), 50 mM NaCl, 3 mM MgCl2, and 300 mM sucrose) and fixed and permeabilised as described above. Before the immunostaining, a treatment with Ambion RNase III (1.2 U, ThermoFisher, AM2290) for 30 min at 37°C was performed, to remove dsRNAs that could interfere with the staining. Cells were blocked for 1 h in blocking buffer (BSA 3% (w/v) in PBS), incubated with the mouse anti-DNA-RNA hybrid S9.6 antibody (1/500, Kerafast ENH001) in blocking buffer overnight at 4ºC, followed by 3 washes with PBS and detection using Alexa Fluor 594 goat anti-mouse secondary antibody (1:800 in blocking buffer) for 1 h at 37°C. Slides were imaged using a Zeiss microscope using Carl Zeiss software. The averages of S9.6 signal intensity per nucleus (A.U., Arbitrary Units) were quantified using the ImageJ software (Fiji).

**Liquid chromatography-tandem mass spectrometry (LC-MS/MS) analysis**

Dried tryptic digests were redissolved in 0.1% TFA by shaking (1200rpm) for 30 min and sonication on an ultrasonic water bath for 10 min, followed by centrifugation (13,000 rpm, 5°C) for 10 min. LC-MS/MS analysis was carried out in technical duplicates and separation was performed using an Ultimate 3000 RSLC nano liquid chromatography system (Thermo Scientific) coupled to a Q-Exactive mass spectrometer (Thermo Scientific) via an EASY spray source (Thermo Scientific). For LC-MS/MS analysis re-dissolved protein digests were injected and loaded onto a trap column (Acclaim PepMap 100 C18, 100µm x 2cm) for desalting and concentration at 8µL/min in 2% acetonitrile, 0.1% TFA. Peptides were then eluted on-line to an analytical column (Acclaim Pepmap RSLC C18, 75µm × 50cm) at a flow rate of 250nL/min. Peptides were separated using a 120-minute gradient, 4-25% of buffer B for 90 min followed by 25-45% buffer B for another 30 min (composition of buffer B – 80% acetonitrile, 0.1% FA) and subsequent column conditioning and equilibration. Eluted peptides were analysed by the mass spectrometer operating in positive polarity using a data-dependent acquisition mode. Ions for fragmentation were determined from an initial MS1 survey scan at 70,000 resolution, followed by HCD (Higher Energy Collision Induced Dissociation) of the top 12 most abundant ions at 17,500 resolution. MS1 and MS2 scan AGC targets were set to 3e6 and 5e4 for maximum injection times of 50 ms and 50 ms respectively. A survey scan m/z range of 400 – 1800 was used, normalised collision energy set to 27%, charge exclusion enabled with unassigned and +1 charge states rejected and a minimal AGC target of 1e3.

**Raw data processing**

Data was processed using the MaxQuant software platform (v1.5.8.3), with database searches carried out by the in-built Andromeda search engine against the Swissprot H.sapiens database (version 20170202, number of entries: 20,183). A reverse decoy search approach was used at a 1% false discovery rate (FDR) for both peptide spectrum matches and protein groups. Search parameters included: maximum missed cleavages set to 2, fixed modification of cysteine carbamidomethylation and variable modifications of methionine oxidation, protein N-terminal acetylation, asparagine deamidation as well as glutamine to pyro-glutamate. Label-free quantification was enabled with an LFQ minimum ratio count of 2. 'Match between runs' function was used with match and alignment time limits of 1 and 20 min respectively.

**Northern blot**

RNA extraction was carried out using RNeasy Mini Kit (Qiagen, 74104) and DNA contaminants were eliminated by in column treatment with DNase I (Qiagen, 79254) according to the manufacturer instructions. RNA (20 µg) was denatured for 10 min at 65°C in sample buffer (formamide 50% (v/v), 2.2 M formaldehyde, 1X MOPS) followed by incubation on ice for 5 min. 10X Dye buffer (glycerol 50% (v/v), Bromophenol Blue 0.3% (w/v), 4 mg/ml ethidium bromide) was added to each sample and they were run on a formaldehyde agarose gel (agarose 0.8% (w/v), formaldehyde 6.5% (v/v) in 1X MOPS) at 5 V/cm in 1X MOPS buffer (0.2M MOPS, 50 mM NaOAc, 10 mM EDTA, RNase-free water). The gel was rinsed twice in water, washed 5 min with denaturation solution (1.5 M NaCl, 0.05 M NaOH), followed by three more washes with 20X SSC before transferring the RNA on an Amersham Hybond N+ membrane (GE Healthcare, RPN303B) overnight using a neutral transfer in 20X SSC. The membrane was UV-crosslinked (Stratalinker, 200 J/cm$^2$ UV) and baked for 45 min at 80ºC. For RNA detection, the membrane was hybridised with a $^{32}$-P labelled TAA(CCCTAA)$_4$probe using UltraHyb-Oligo solution (ThermoFisher #AM8663) at 42ºC overnight, followed by 10 min wash with 2XSSC/0.1% (w/v) SDS, 2 min wash with 0.2XSSC/0.1% (w/v)

SDS at 42°C and a final 5 min wash with 2XSSC/0.1% (w/v) SDS solution. Radioactive signals were detected by phosphor-imager (Amersham Biosciences).

**RNA dot blot**

RNA (5 µg) was incubated in 0.2 M NaOH, denatured at 65°C for 10 min and incubated on ice for 5 min. Finally, RNA was spotted onto a positively charged Amersham Hybond N+ membrane (GE Healthcare, RPN303B) using a dot-blot apparatus. Membranes were UV-crosslinked (Stratalinker, 200 J/cm$^2$ UV), heated for 30 min at 80°C and hybridised overnight at 42°C with the following digoxigenin (DIG)-labeled probes: telomeric C-rich oligonucleotide TAA$(CCCTAA)_4$; telomeric G-rich probe ATT$(GGGATT)_4$ and 18s rRNA probe CCATCCAATCGGTAGTAGCG. The probes were prepared using 3' end labelling kit (Roche, 03353575910) according to the manufacturer's instructions and diluted in hybridisation buffer (6X SSC, 0.1% SDS, 1% milk). The membrane was then washed 2 x 5 min with pre-warmed 2X SSC/0.1% SDS and 1 x 2 min with pre-warmed 0.2X SSC/ 0.1% SDS, then submerged in 2X SSC for 5 min and washed in Tween-20 in maleic acid 0.3% (v/v) for 5 min. Signal was detected using the anti-DIG-alkaline phosphatase antibodies (1:20000, Roche, 11093274910) and CDP-Star (Roche, 11685627001) following the manufacturer's instructions. Images were captured using the Amersham Imager 680 (GE Healthcare) and analysed using the Image Studio Lite software. 18s rRNA signal was used for normalisation.

**Additional DNA-RNA Immunoprecipitation (DRIP) experiments**

Additional DRIP experiments were performed using sonicated nucleic acids (Arora et al. 2014). Samples were sonicated using Bioruptor (Diagenode) to obtain around 500 bp long fragments and 30 µg of digested nucleic acids were treated or not with 10 µl of RNase H overnight at 37ºC. 5 µg of sample was incubated with complexes of 10 µl of the S9.6 antibody (Kerafast ENH001) and 30 µl of Dynabeads Protein A for 2 h at 4ºC. 1 µg of sample was used as input.

**ATP hydrolysis assay**

The ATP hydrolysis reaction (10 µl) contained 20 mM Tris-HCl pH 7.5, 4 mM magnesium acetate, 1 mM DTT, 0.5 M ATP and 3 nM radioactive ATP $^{32}$P (specific activity 6000 Ci/mmol). The reaction was initiated by adding indicated amounts of SKI protein complex, incubated for 3.5 hrs at 37ºC before stopping the reaction with 50 mM EDTA. 1 µl of each reaction was spotted on TLC plates (Thin layer chromatography, PEI cellulose, Merck). The plates were dried and released radioactive phosphate (Pi) was separated from ATP by chromatography in a solution containing 0.15 M lithium chloride and 0.15 M formic acid. The plates were again dried before exposing to phosphor screens. The screens were scanned using FLA-5100, the free 'Pi' signal was quantitated using image J (Fiji).

**Preparation of DNA/RNA substrates**

The sequences of all oligonucleotides and substrates used are listed in Tables S1 and S2. The RNA and DNA oligonucleotides were synthesized and PAGE purified by SIGMA. The oligonucleotides were labelled with radioactive $^{32}P$ at 5' end by T4 polynucleotide kinase. The labelled oligo was annealed to the complementary non-labelled oligo in a 1:2 ratio in polynucleotide kinase buffer, heated at 95ºC for 5 min followed by cooling overnight.
